# Unveiling Gene Perturbation Effects through Gene Regulatory Networks Inference from single-cell transcriptomic data

**DOI:** 10.1101/2024.05.10.593314

**Authors:** Clelia Corridori, Merrit Romeike, Giorgio Nicoletti, Christa Buecker, Samir Suweis, Sandro Azaele, Graziano Martello

## Abstract

Physiological and pathological processes are governed by a network of genes called gene regulatory networks (GRNs). By reconstructing GRNs, we can accurately model how cells behave in their natural state and predict how genetic changes will affect them. Transcriptomic data of single cells are now available for a wide range of cellular processes in multiple species. Thus, a method building predictive GRNs from single-cell RNA sequencing (scRNA-seq) data, without any additional prior knowledge, could have a great impact on our understanding of biological processes and the genes playing a key role in them. To this aim, we developed IGNITE (Inference of Gene Networks using Inverse kinetic Theory and Experiments), an unsupervised machine learning framework designed to infer directed, weighted, and signed GRNs directly from unperturbed single-cell RNA sequencing data. IGNITE uses the GRNs to generate gene expression data upon single and multiple genetic perturbations. IGNITE is based on the inverse problem for a kinetic Ising model, a model from statistical physics that has been successfully applied to biological networks. We tested IGNITE on murine pluripotent stem cells (PSCs) transitioning from the naïve to formative states. Using as input only scRNA-seq data of unperturbed PSCs, IGNITE simulated single and triple gene knockouts. Comparison with experimental data revealed high accuracy, up to 74%, outperforming currently available methods. In sum, IGNITE identifies predictive GRNs from scRNA-seq data without additional prior knowledge and faithfully simulates single and multiple gene perturbations. Applications of IGNITE range from studying cell differentiation to identifying genes specifically active under pathological conditions.

## 2 Introduction

The correct functioning of cellular processes requires regulated interactions of several components. By studying each individual element separately, we cannot grasp how the organisation of the individual components involved in a specific process can lead to complex behaviours. However, studying the whole system is still a challenging task. Among all the elementary components of the cell, genes are the key necessary elements in regulating biological processes by interacting and organising themselves into Gene Regulatory Networks (GRN). To study the interactions between genes and how they shape the cell, it is possible to experimentally perturb the regulation of individual genes, forcing their activation or deactivation. An experimental technique to perturb the GRN by deactivating specific genes is the knockout. This technique allows to measure the effects of individual or multiple gene deactivations on the regulation of other genes, and how this affects phenomenological cell dynamics. However, given the multitude of genes on which it is possible to focus, an experimental approach would be too time-intensive and costly. In silico models represent a viable alternative, as they can infer complex gene interactions and predict the outcome of genetic perturbation, ultimately revealing key regulatory mechanisms.

Emerging computational strategies aim to predict how perturbations affect cell phenotypes. To effectively apply these methods, it is essential to understand their capabilities and limitations. For instance, the reliance on data obtained from experimental perturbations for the training step limits the applicability of these models [1]. Furthermore, numerous approaches necessitate the integration of multi-omics data as input knowledge to the algorithm. This implies having access to these data by performing different measurements on the same biological system. Alternatively, some methods can infer GRN interactions only from unperturbed single-cell RNA sequencing (scRNA-seq) data [2]. However, they focus only on some specific aspects, such as GRN inference, or generation of wild-type data, or perturbation simulations.

Among the approaches for GRN inference, RE:IN [3, 4, 5, 6] is capable of inferring interactions among numerous genes using a Boolean network modelling approach and simulating knockouts of one or two genes. However, RE:IN relied on both knockout experiments and bulk transcriptomic data, lacking single-cell resolution, and needed to include other types of data (ChIP sequencing data, named ChIP-seq, and literature-curated interactions) to constrain the model. Moreover, RE:IN should explore a number of possible networks that increase exponentially with the number of genes, making the inference task computationally demanding. Another method of inferring the GRNs is MaxEnt. It employs the maximum entropy principle to infer genetic interaction networks from gene expression patterns [7]. The GNR is obtained by maximising the entropy while satisfying the specific constraints given by the data [8, 9]. However, MaxEnt can infer only undirected interactions, and it is not suitable for generating perturbed data. SCODE [10] is one of the gold standard models for inferring directed weighted networks starting from scRNA-seq data and exploiting pseudotime. It describes gene expression dynamics with a set of coupled linear ordinary differential equations. Moreover, it can generate new data, but it has not been validated for the simulation of gene perturbation. CellOracle [11] is a network-based machine learning framework. It uses scRNA-seq data from different clusters of cells and map active regulatory connections onto a GRN framework computed using scATAC-sec data (or ATAC-seq data). It produces GRNs for each cluster of the dataset and stimulates the effects of perturbing a transcription factor with a known binding motif. CellOracle focuses on the role of GRN perturbation in modifying cell phenotypes, without describing how GRN can generate unperturbed phenotypes.

All these approaches study only some specific features that resemble the real system, which we summarised in Table 1. However, it would be essential to develop interpretable models that require only one type of unperturbed data (e.g., single cell transcriptomics), simplifying algorithm usage and providing a description that includes both GRN inference, a generative rule explaining how GRN can produce cell phenotypes, with the capacity to simulate the effects of gene perturbations.

**Table 1:** Comparison of different computational methods for GRN inference, highlighting their input data type, requirement of prior information, capabilities of generating new data, implementation of multiple gene knockouts (KO) simulations, and ability to infer directed interactions. ^*^These features are theoretically possible, but were not implemented

| Methods | Input data resolution | Prior information | Generative model | Multiple KO | Directed |
| --- | --- | --- | --- | --- | --- |
| RE:IN | Bulk RNA-seq | ChIP-seq data and literature-curated interactions | Yes | Yes | Yes |
| MaxEnt | Bulk or scRNA-seq | Only transcriptomics | No* | No | No |
| SCODE | scRNA-seq | Only transcriptomics | Yes | No* | Yes |
| CellOracle | scRNA-seq | scATACseq or ATAC-seq | No | No* | Yes |
| IGNITE | scRNA-seq | Only transcriptomics | Yes | Yes | Yes |

**Table 2:** Fraction of Agreement for different KO simulations with the three methods considered: IGNITE, SCODE [10] and CellOracle [11]. Instances where the values are lower than what would be expected randomly are highlighted with ^*^ (see Methods for details).

|  | **Rbpj** | **Etv5** | **Tcf7l1** | **Triple** |
| --- | --- | --- | --- | --- |
| IGNITE | 0.65 | 0.74 | 0.30* | 0.67 |
| SCODE | 0.30* | 0.22* | 0.30* | 0.48* |
| Cell Oracle | 0.39* | 0.30* | 0.13* | 0.62 |

In response to these modelling challenges, we propose IGNITE (Inference of Gene Networks using Inverse kinetic Theory and Experiments). It is an unsupervised computational approach, based on concepts of statistical physics, that starts from unperturbed scRNA-seq data to infer GRN models capable of simulating unperturbed or perturbed cell phenotypes. In particular, it relies on the inverse problem for a kinetic Ising model [12], following the approach proposed by Zeng et al. in [13]. The Ising model was initially developed to describe ferromagnetism and has found applications in modelling several many-body systems, including biological networks and neural networks [7]. Here, we leverage its versatility to develop a computational model describing the GNR by exploiting all the features reported in Tab. 1.

We applied IGNITE to the study of the developmental progression of pluripotent stem cells (PSCs). Pluripotency is the potential to give rise to all cells found in the adult and is observed in the naive epiblast of the preimplantation embryo. Upon embryo implantation, epiblast transition to a different pluripotency phase, termed formative phase, characterised by distinct gene expression profiles, epigentic state and developmental features. During this transition, naive pluripotency markers are down-regulated, epigenetic modifiers and formative markers are activated, and cells become competent in the formation of the germ layer and the specification of primordial germ cells (E6.5–7.5) [6]. Pluripotent cells of the early embryo can be captured in vitro as naive Embryonic Stem cells (ESC), which retain pluripotency and the capacity to progress from the naive to the formative state. Several genetic approaches have been used to identify genes crucial for the maintenance of the naive state, or the transition to the formative state. However, a global understanding of how the transition is regulated is still missing.

Previous approaches focused mainly on characterising the state of naive pluripotency and elucidating the functional interactions among the molecular components that govern and sustain it [14, 15, 16, 17]. Moreover, efforts focused on modelling the interactions of these core genes and their dynamics under different cell culture conditions. For instance, Walker et al. [14] experimentally deduced the possible GRNs of mouse ESCs in the pluripotency state, while other approaches, relying on established GRNs, characterised the dynamics of a few interacting genes using differential equations [15, 16, 17]. However, while informative and precise, these approaches were limited to describing the interactions of a limited number of genes.

The Boolean network approach has been applied not only to bulk RNA-seq data (as done with RE:IN), but also to single-cell data [18, 19]. These methods allowed to have single cell resolution data, but still required the integration of different types of data, e.g. ChIP-Seq and knockout experiments for [18] and qRT-PCR and single-molecule-mRNA fluorescent in situ hybridisation (sm-mRNA-FISH) in [19]. Other approaches focused only on transcriptomics data (qRT-PCR and scRNA-seq) [20, 21, 22]. However, a common issue is the fact that these approaches should explore a number of possible networks that increase exponentially with the number of genes, making the inference task computationally demanding.

Other computational strategies, including tree-based models, correlation and partial correlation analysis, information theory, and differential equations, have been used to model differentiation from unperturbed single-cell multi-omics data [2, 11]. However, even in studies on differentiation, these methods focus primarily on inferring GRNs, as previously highlighted, without elucidating how these GRNs can generate differentiating cells or inspecting the role of perturbations in differentiation.

Hence, it is essential to study this system to have a reliable model capable of generating all the observed cell phenotypes and being able to simulate knockout to investigate gene function during the transition. Starting from a novel 10X scRNA-seq dataset of murine ESCs progressing from naive to formative state, IGNITE inferred the GRN. Using this obtained GRN, IGNITE can generate new synthetic data without depending on preexisting biological knowledge, such as different data types (e.g., ATACseq data) or literature-based information (e.g., established interactions). Remarkably, the generated wild-type data were statistically comparable with the experimental measurements. IGNITE also simulated the knockout of three genes, Rbpj, Etv5, and Tcf7l1, both individually and concurrently perturbing all three genes simultaneously. Strikingly, these simulations were largely supported by experimental data. Of note, Tcf7l1 is a transcription factor regulated at the level of protein stability, rather than transcriptionally. Simulations of Tcf7l1 inactivation by IGNITE and other methods were not accurate, revealing limitations of approaches based only on transcriptomic data.

IGNITE represents a model-based computational approach to generate new single-cell phenotypes as patterns of (de)activation of genes and to simulate perturbation experiments thanks to the inference of the GRN describing the biological system studied, outperforming current gold standard methods ([10, 11]).

## 3 Results

### 3.1 Deciphering Pluripotent Stem Cell Dynamics through scRNA-seq

The IGNITE algorithm comprises three key steps (Fig. 1A). (i) **Processing**: scRNA-seq input data undergo Quality Control and logarithmic normalisation, followed by pseudotime (PST) computation and application of the Mini-bulk (MB) approach (for further details, refer to the Methods section). The PST computation requires data dimensionality reduction and clustering. The PST algorithm Slingshot [23] orders individual cells along a pseudotime trajectory, assigning to each cell a pseudotime value that represents its position along that trajectory [24]. The pseudotime implementation allows us to leverage the variability among cells to obtain a dataset where individual cells are aligned along a pseudotime trajectory from a dataset that contains only a few time points. After PST, we compute the moving average of different subsets of cells, thus generating “Mini-Bulks” (MB) (see Methods for details). The MB approach is particularly useful to address the problem of having high false negative dropout rates, typical of scRNA-seq data[25]. Applying PST and MB strategy ensures the averaging of similar cells along the differentiation trajectory. The algorithm models gene expression as on/off spins states within a kinetic Ising model framework, solving the inverse Ising problem to infer gene interactions [13]. To do so, IGNITE computes gene activity (GA). This consists of assigning state +1 to active genes (high gene expression) and -1 to inactive genes (low gene expression) (see Methods for details). (ii) **Modelling**: IGNITE infers the GRN model based on GA. Consequently, genes are modelled as spins of a kinetic Ising model with unknown pairwise couplings between them [13, 7]. The system dynamical rule follows the Glauber dynamics with asynchronous spin updates [12]. IGNITE solves the inverse Ising problem by maximising the likelihood of the system. In doing so, IGNITE generates multiple GRNs, and, among them, selects the one with the lowest distance between input and generated data correlation matrices. (iii) **Data Generation**: using the GRN, IGNITE generates new data under desired conditions, either wild-type (WT) or upon gene perturbations (knockout, KO). The IGNITE output includes (1) the inferred effective GRN, (2) the generated gene activity data under wild-type (it replicates the system condition of the original data), and (3) the effects of specific gene perturbations, like knockouts, on the gene activity (see Methods for details).

**Figure 1.**
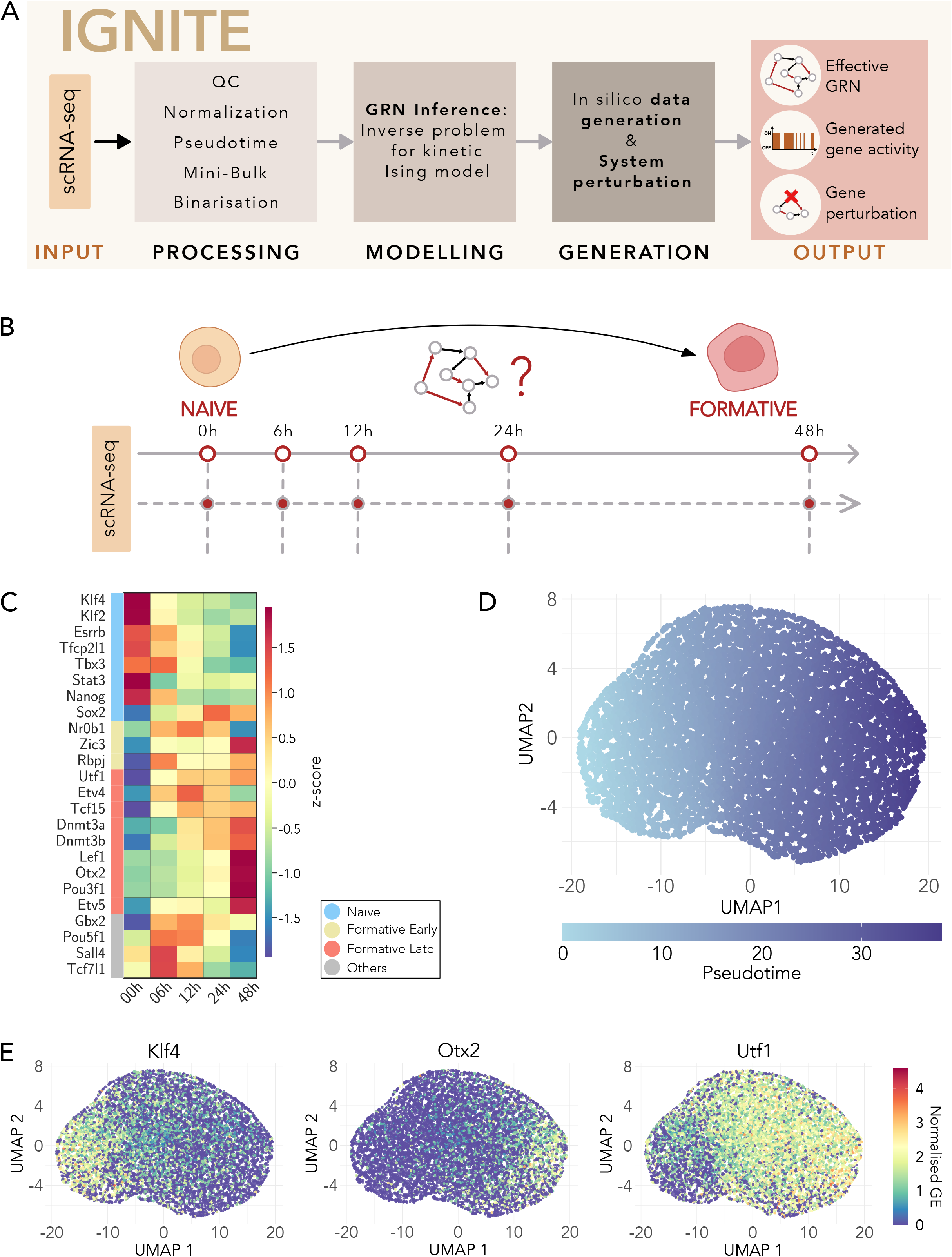
A. Overview of the IGNITE workflow: from the scRNA-seq input, through processing, modelling, and data generation, to the final outputs: (i) effective inferred Gene regulatory Network(GRN), (ii) wild-type generated gene activity data, and (iii) gene activity generated using inferred GRNs with gene perturbations. QC in dicates Quality Control. B. Studied biological system: temporal dynamics of PSCs differentiation, encompassing the transition through distinct stages from naive to formative. The considered transcriptomic dataset is 10X scRNA-seq starting in 2i+LIF (2iL). The points represent 5 time points at which specific cells were sampled and measured after the removal of 2iL. C. Average gene expression z-score of the log-normalised scRNA-seq input dataset. For each gene, the average is computed for all cells that belong to the same time point. We considered 24 genes, divided into four different gene categories, naive (light blue), formative early (yellow), formative late (red), and others (grey). We classified these genes following Carbognin et al. [6]. D. Two-dimensional UMAP visualisation of the log-normalised scRNA-seq dataset, with each point representing an individual cell. The gradient colour scale corresponds to the pseudotime value of each cell, indicating the time progression within the dataset. The pseudotime ranges from the value of 0 to the final value of 38.22. UMAP1 and UMAP2 indicate the two dimensions of the UMAP space. E. Visualisation of two-dimensional UMAPs of the log-normalised scRNA-seq dataset, showing the expression patterns of three key genes: Klf4, Otx2, and Utf1. Each point represents an individual cell coloured according to the normalised gene expression (GE) level of the respective gene. UMAP1 and UMAP2 indicate the two dimensions of the UMAP space.

As our biological system of interest, we selected differentiating PSCs. Our PSC cultures were maintained in a medium containing LIF and 2i (2iL). This chemically defined culture condition ensures the naive pluripotent state of stem cells [26, 27]. When the 2iL medium is removed, the PSCs undergo a series of developmental stages, starting from an initial state, naive, progressing to the formative state (Fig. 1B) [28]. We removed the 2iL medium and measured gene expression at different time points using scRNA-seq measures to have information on the dynamics of gene activity during differentiation. We focused on 24 genes known in the literature for their driving role in the decision-making process of differentiating PSCs [3, 6, 29, 30]. We concentrated on the first 48 hours of PSC development, as we want to focus on inferring the interactions that guide the transition between naive and formative cells, analysing a total of 9894 cells.

During this transition, we observed four main patterns of behaviour (Fig. 1C): a first group of genes, named “naive”, exhibit higher activity in the naive state of the cell, which subsequently decreases. The “formative early” genes have low gene expression at the initial time point, and then they are activated at the immediately subsequent time points, while the “formative late” genes are activated at later steps. Lastly, a mixed group of genes display variable activity at each step, without a specific trend. This behaviour is in agreement with previously reported bulk transcriptomics data [6] (Fig. S1A). To compute the PST, we projected the data in a lower-dimensional space using t-distributed Stochastic Neighbour Embedding (t-SNE) [31] and Uniform Manifold Approximation and Projection (UMAP) [32] (Fig. S1B) and we clustered the data (Fig. S1C), see Methods for details. To validate the biological significance of the obtained clusters, we investigated the distribution of gene expression in the genes within each cluster (refer to Fig. S1D for three examples). In addition, we excluded cluster $1$ from subsequent analysis due to its predominant composition of 2Cell-like cells (2CLCs), identified with confidence through a comprehensive analysis of differential gene expression [33]. We found as up-regulated markers genes such as Eif1a-like genes and Zscan4. This cluster was omitted from further investigation as it diverged from the main scope of our analysis, investigate the transition from formative to naive state. After computing the PST, we checked the PST values distribution for each time point (Fig. S1E), observing agreement between sampling time and PST ordering. We verified that the expected temporal expression patterns of the genes were maintained even after PST ordering. In the low-dimensional UMAP space (Fig. 1D), it is possible to observe the PST assigned to each cell. Cells with lower PST exhibit higher gene expression of naive genes (e.g. Klf4, in Fig. 1E) compared to formative genes. Conversely, formative genes displayed increased activity in cells with higher pseudotime values, as expected (e.g. Otx2 and Utf1 in Fig. 1E). Gene expression dynamics, after PST and MB processing, is preserved. Indeed, its behaviour over time (Fig. S1F) is analogous to that described in Fig. 1C. Finally, we calculated GA and still observed the expected transitions from the naive to the formative states (Fig. S1G).

### 3.2 GRN inference and WT data generation: IGNITE and SCODE

IGNITE uses only scRNA-seq data as input to infer a set of 250 GRNs (Fig. 1A). It employs the correlation matrix of the generated data and its time derivative to infer the most accurate and effective GRN. It achieves this by comparing the correlation matrices between the input and generated datasets, quantifying their difference by means of their Euclidean distance. We defined this quantity as Correlation Matrices Distance, CMD (see Methods for details). The GRN with the lowest CMD (0.43) (Fig. 2A), which best reproduces the initial data correlations, is selected. Statistical validation by a two-sample t-test yielded a t-statistic *t* = -43.49 and a p-value *p* = 1.04 × 10^−171^, confirming a significant difference between the IGNITE model and the null model. We compared the input experimental data (processed scRNA-seq data with PST and MB) with those generated by the IGNITE-selected GRN and found them qualitatively congruent in terms of correlation (Fig. 2B and 2C, respectively). Furthermore, we evaluated the quality of IGNITE-generated data by clustering the original and generated binarised datasets. In Fig. 2D, the heatmap illustrates the gene expression patterns within the clusters of the input dataset. In the first cluster on the left, naive genes are active, while formative genes are largely inactive (naive cluster). Progressing to the right, there is a clear shift as naive genes become less expressed and formative genes predominate (formative cluster). As expected, the others group remained consistently active throughout the simulation. Comparing the input data with the IGNITE generated ones (Fig. 2E) we verified that the two clusters had a similar representation and composition. In the input dataset, the naive/formative cluster contains 26% / 74% of the cells, and in the generated dataset it contains 20%/80%. Principal component analysis (PCA) on experimental (Fig. 2F) and simulated (Fig. 2G) data further enhanced the high similarity.

**Figure 2.**
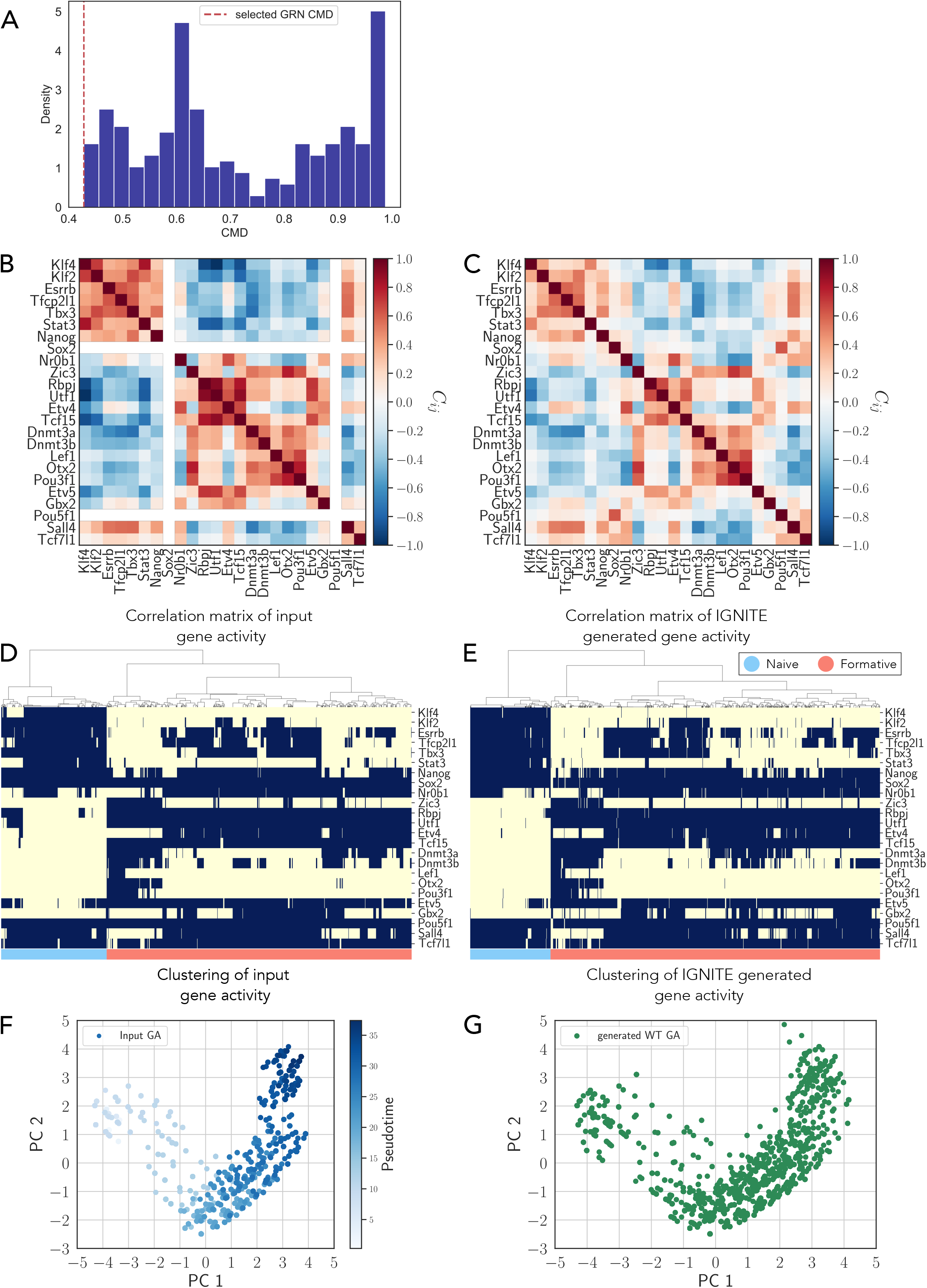
A.Density plot of the Correlation Matrices Distance (CMD) values for the 250 inferred GRNs. The red dashed line represents the selected GRN with the lowest CMD. B. Pearson correlation matrix for the gene activity of the input dataset (scRNA-seq data with LogNorm, PST, and MB). C. Pearson Correlation matrix for IGNITE-generated gene activity dataset. D. Heatmap with hierarchical clustering of gene activity of the input dataset. The clustering algorithm used is Ward’s method. Each row represents a gene, while each column corresponds to an individual cell. The color indicates inactive (−1, yellow) or active (+1, blue) gene activity. The dataset has 9547 cells. E. Heatmap with hierarchical clustering of gene activity of IGNITE-generated data. Methodology for visualisation as in Fig. 2D. 9547 cells were simulated. F. Two-dimensional Principal Component Analysis (PCA) scatter plot representing the gene activity, GA, for the input dataset (scRNA-seq data with LogNorm, PST, and MB). Each point corresponds to a single cell, and the colour intensity reflects the pseudotime value of the cell. PC1 and PC2 indicate the two dimensions of the PCA space. G. Two-dimensional PCA scatter plot representing the IGNITE-generated wild-type gene activity data, WT GA, using the same dimensional reduction approach as in Fig. 2F. PC1 and PC2 indicate the two dimensions of the PCA space.

Next, we compared the results obtained with IGNITE against those generated by SCODE. To infer the GRN with SCODE, we searched for the optimal parameters as suggested by the authors in [10] (Fig. S2A and S2B, see Methods for details).

Looking at the clustering of the SCODE gene expression data we observed that there are similarities between the input and the generated data (Fig. S2C and S2D, respectively), with elevated expression of some naive genes (e.g. Klf2/4) in naive cells, and of formative genes Rbpj and Utf1 in formative cells. The correlation matrices of input and SCODE generated data are quantitatively comparable (Fig. S2E and S2F), revealing an agreement between the input and generated data, as shown also by the PCA (Fig. S2G). Analysis of the correlation matrices revealed a CMD of 0.51. This value is statistically relevant (z*s*co*r*e = −428.30), but higher than the one obtained by IGNITE. We conclude that IGNITE displays generative capacity comparable to the benchmark model SCODE.

### 3.3 Simulating Gene Knockouts in Pluripotent Stem Cell with IGNITE

To assess the predictive capabilities of IGNITE, we simulated the knockout Rbpj, Etv5, and Tcf7l1 genes individually (single KOs) as well as the knockout of these genes simultaneously (triple KO) by removing their interactions from the effective GRN. Using 4 different KO GRNs, we generated the knockout datasets (see Methods for details).

Studying the generated KO data, we observed whether the gene activity of the remaining genes increased or decreased compared to the wild-type by measuring the difference between the average activity per gene for KO and WT simulated data. To quantify these changes, we calculated the difference in gene activity/expression between KO and WT, scaled to the highest absolute value (scaled KO-WT difference, Fig. 3A and Fig. 3B). We then compared the scaled KO-WT difference of generated data with the one obtained from experimental data. This last quantity is obtained by scaling the experimental log2FC data for single KO data [30] and triple KO data [29]. Using these scaled KO-WT difference values we calculated the Fraction of Agreement (FoA), which measures the fraction of genes for which generated and experimental gene expression changes are in the same direction (see Methods for details) (Tab. 1). Single KO simulations are characterised by increased activity for naive genes and reduced for formatives, in agreement with previous reports [30, 29]. The FoA for Rbpj and Etv5 KO is equal to 0.65 and 0.78, respectively. Both values are statistically significant compared to null models. On the contrary, the KO of Tcf7l1 could not be faithfully predicted, an expected result as Tcf7l1 is known to be regulated at the level of protein stability and DNA affinity, which cannot be captured by scRNA-seq [34].

**Figure 3.**
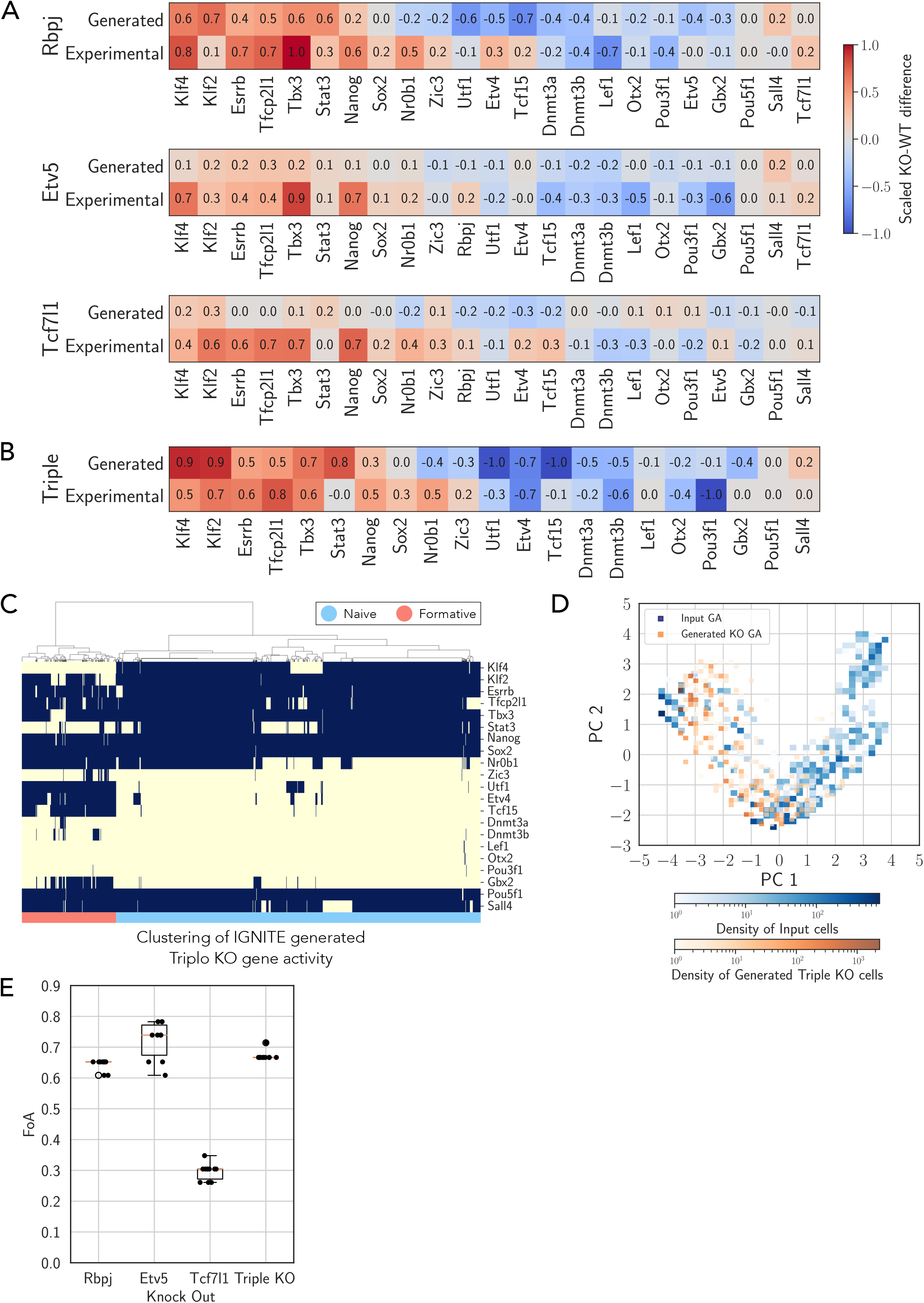
A. Scaled KO-WT difference for Rbpj, Etv5 and Tcf7l1 simulations with IGNITE and comparison with experimental scaled KO-WT difference data [30]. For each gene, to compute the simulated scaled KO-WT difference, we measured the scaled difference between the average fraction of active cells under wild-type and knockout conditions. For the experimental data, we scaled the log2FC values measured in [30]. The scaling of all quantities is between -1 and +1 to facilitate the comparison between different datasets (see Methods for details). B. Scaled KO-WT difference for Triple KO simulation with IGNITE and comparison with experimental scaled KO-WT difference [29]. Methodology for computing the simulated end experimental scaled KO-WT difference as in Fig. 3A (see Methods for details). C. Heatmap with hierarchical clustering of the IGNITE-generated triple KO GA. Methodology for visualisation as in Fig. 2D. 9547 cells were simulated. D. Two-dimensional PCA scatter plot representing the gene activity, GA, of the input dataset (scRNA-seq data with LogNorm, PST, and MB) and the IGNITE-generated triple KO GA. The colour gradient represents the cell density within each square area, with separate scales for the input (blue) and the generated triple KO cells (orange). PC1 and PC2 indicate the two dimensions of the PCA space. E. Whisker plot of the Fraction of Agreement, FoA, values for each type of KO (Rbpj, Etv5, Tcf7l1 and all three combined) for 10 GRN (inferred with IGNITE) that exhibited the lowest CMD values (selected from 250 tested models). The box is drawn to represent the lower and upper quartiles, with an orange line indicating the median. The whiskers extend from the box to illustrate the full range of the non-outlier data. Any data points beyond the whiskers are referred to as outliers.

For the triple knockout of Rbpj, Etv5, and Tcf7l1, both the simulations and experimental data displayed a strong increase in naive genes and a decrease in formative genes, with a significant FoA of 0.67. Triple knockout of Rbpj, Etv5, and Tcf7l1 has been reported to expand for several passages under conditions promoting differentiation, while retaining expression of naive markers in the vast majority of cells [29]. As IGNITE could simulate the cellular composition of wild-type differentiating cells (Fig. 2E), we asked whether it could simulate the cell composition of triple KO cells (Fig. 3C). Strikingly, most cells (79%) displayed activation of all naive genes, the remaining 21% exhibited activity of some formative genes (Utf1, Etv4 and Tcf15) and inactivation of only two naive genes (Klf4, Stat3).

Comparing the PCA of the triple KO gene activity generated with IGNITE with the PCA of gene activity input data with MB (Fig. 3D) we observed that the triple KO IGNITE data closely mirror the gene activity patterns observed at the initial time points of the input data, which are marked by a predominance of naive state cells. This result highlights IGNITE ability to accurately replicate the expected naive state gene activity in the triple KO, consistent with established literature findings [29].

To evaluate the results of IGNITE we simulated the single and triple gene perturbation with SCODE and CellOracle [11]. We computed the scaled KO-WT difference for SCODE and CellOracle simulations, and compared it with the experimental log2FC (Fig S3A and S3B. Qualitatively, the results obtained did not show a clear resemblance to the experimental findings. Measuring quantitatively the performance of all considered methods, IGNITE outperformed the others in gene perturbation simulations. IGNITE had a higher FoA value for Rbpj, Etv5, and triple gene knockout compared to SCODE and CellOracle 2. Furthermore, SCODE generated data that were not comparable to those in the early stages of differentiation under the WT condition (Fig. S4A). Similarly, in the PCA space, the data simulated with CellOracle predominantly overlapped with regions characterised by low levels of gene expression of naive and formative genes of wild-type data and only partially overlapped with areas where naive gene expression was highest (Fig. S4B and S4C). On the contrary, IGNITE successfully predicted the expected cell composition in the data generated after triple KO, thus forecasting the effects of gene perturbations on PSCs.

For the results obtained so far with IGNITE, we selected the effective GRN as the one with the lowest CMD (Fig. 1A). As a further analysis, we verified that the IGNITE performances in simulating perturbations in the biological system are stable for any GRNs with low normalised distances. For this reason, we selected the 10 GRNs with the lowest CMD values and calculated the FoA values for single and triple knockouts (Fig. 3E). FoA values are very stable, suggesting that IGNITE is robust to variations within the top-performing GRN models. This robustness ensures that the predictions made by IGNITE are not artefacts of a single GRN but are representative of underlying biological processes.

### 3.4 Assessing the Accuracy of GRN IGNITE Inferences Against Experimental Benchmarks and other Inference Methods

Next, we analysed the GRN inferred with IGNITE, asking whether it contained gene interactions previously validated using independent techniques. The GRNs were visualised as interaction matrices in which the interaction from gene *j* (columns) to gene *i* (rows) is denoted by the corresponding entry (*i, j*) in the matrix. Notably, positive interactions are prevalent within the naive group and within the formative group, as previously reported. A predominance of negative interactions is observed between the naive and formative genes (Fig. 4A).

**Figure 4.**
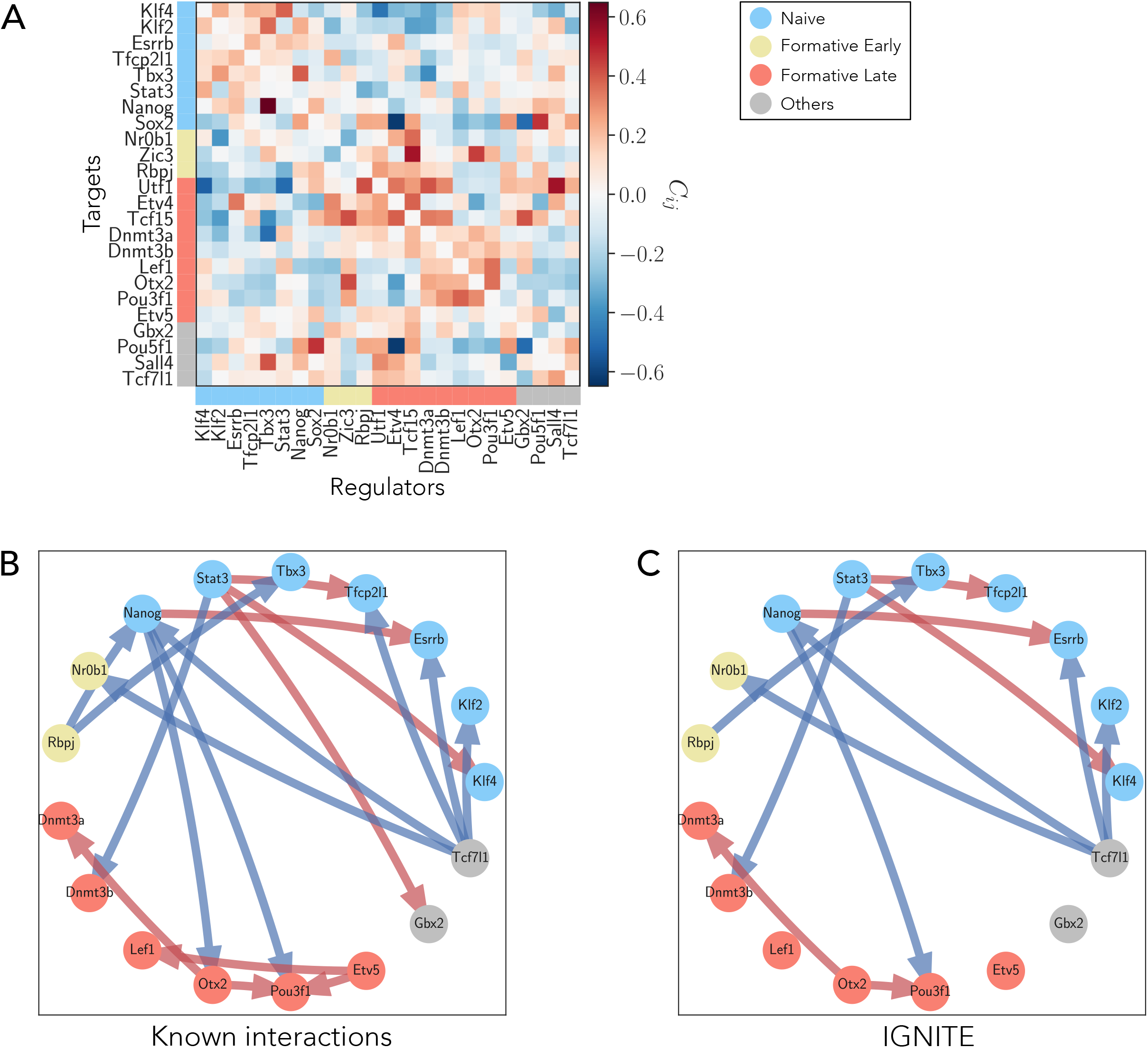
A. IGNITE GRN interaction matrix inferred using the input dataset (scRNA-seq data with LogNorm, PST, and MB). The genes are grouped as in Fig. 1C: naive (light blue), formative early (yellow), formative late (red), and others (grey). The interaction values can be negative (blue) or positive (red). The colour of each matrix element (*i, j*) represents the intensity of the interaction value from gene *j* to gene *i*. B. GRNs of the set of 18 interactions known from the literature. Only the genes involved in these interactions are shown as nodes. Genes are coloured by their group type as in Fig. 4A. The red arrows indicate positive (activating) interactions, and blue arrows negative (inhibiting) interactions, respectively. C. GRN of the subset of interactions known from literature correctly inferred with IGNITE. The arrows represent the interactions that IGNITE inferred with the correct sign.

For comparison, we inferred the GRN with three additional methods, SCODE, CellOracle and MaxEnt. The MaxEnt GRN interaction matrix shows positive interactions among naive genes, and also among some formative genes, as expected (Fig. S5A). However, the pattern of negative interactions between naive and formative groups observed in the IGNITE GRN is not clearly distinguishable. We computed the interaction matrix also using SCODE and CellOracle (Fig. S5B and S5C, respectively). The SCODE matrix does not show recognisable interaction patterns, as there are no distinct blocks of positive or negative interactions. The matrix shows fewer strong interactions. CellOracle inferred five GRNs, one for each cluster identified in the input scRNA-seq dataset. These CellOracle GRNs present interaction patterns similar to IGNITE, with positive interactions within the naive genes and the formative genes. Only the interaction strength changes among the clusters, while the patterns are preserved.

To quantitatively compare the inferred GRNs, we have identified 18 experimentally validated interactions from the literature (Fig. 4B) [29, 35, 36, 37, 38, 39, 40]. We considered an interaction as experimentally validated when a factor has been shown to directly interact with the promoter of target genes, and its genetic inactivation or over-activation results in a significant change in the levels of its direct targets. In Fig. 4C, we presented the IGNITE subnetwork, which includes only the genes involved in the known interactions. Its links are the known interactions correctly inferred. Importantly, statistical tests on null models confirmed that these interactions cannot be attributed to random chance (Fig S6A).

From the known interactions, we calculated the fraction of correctly inferred interactions (FCI) as the fraction of all inferred interactions that have the same sign as the known interactions (Tab. 3) (see Methods for details). We focused exclusively on the signs of the interactions since their strength depends on the model and remains experimentally unmeasurable. Given the undirected nature of the MaxEnt GRN, we did not consider the directionality of the known interactions for this method. The FCI for all methods was comparable, indicating a similar ability to identify interactions among all methods tested.

**Table 3:** Fractions of known interactions correctly inferred, FCI, for the 4 considered inference methods using as input the scRNA-seq dataset log-normalised logNorm and processed with PST and MB.

| Metodo | FCI |
| --- | --- |
| IGNITE | 0.67 |
| MaxEnt | 0.72 |
| SCODE | 0.67 |
| CELLORACLE | $0.67 \pm 0.4$ |

### 3.5 Integrating Experimentally Validated Interactions into IGNITE

IGNITE takes as input only scRNA-seq data, and does not require prior knowledge (Fig. 1A), as it selects GRN by minimising the CMD. However, other approaches integrate different experimental datasets and use them as prior knowledge. For example, CellOracle incorporates interactions matrices inferred from ATAC-seq data. We therefore asked whether using experimentally validated interactions as prior knowledge would improve the performance of IGNITE. To do so, we searched for GRNs that maximised FCI. We identified a set of ten GRNs containing 14 of 18 experimentally validated interactions (FCI=0.78).

To investigate the relationship between the two approaches, we calculated the CMD and FCI for all GRNs generated by IGNITE (Fig. 5A). The 10 GRNs maximising the FCI (orange dots) are not the same as minimising the CMD (red dots) that we analysed in Fig. 3E. However, the two parameters anticorrelate (Spearman correlation *ρ* = −0.69, with *p*-value=5.82*e* − 36), potentially indicating that the two approaches might identify GRNs with similar properties. We performed KO perturbation analyses for those 10 GRNs maximising the FCI and obtained highly variable results, with some GRNs displaying FoA values comparable to a null model (Fig. 5B). This is in stark contrast to the performance of the 10 GRNs selected minimising CMD, without prior knowledge (Fig. 3E). Thus, we conclude that using experimentally validated gene interactions as prior knowledge does not improve IGNITE performance.

**Figure 5.**
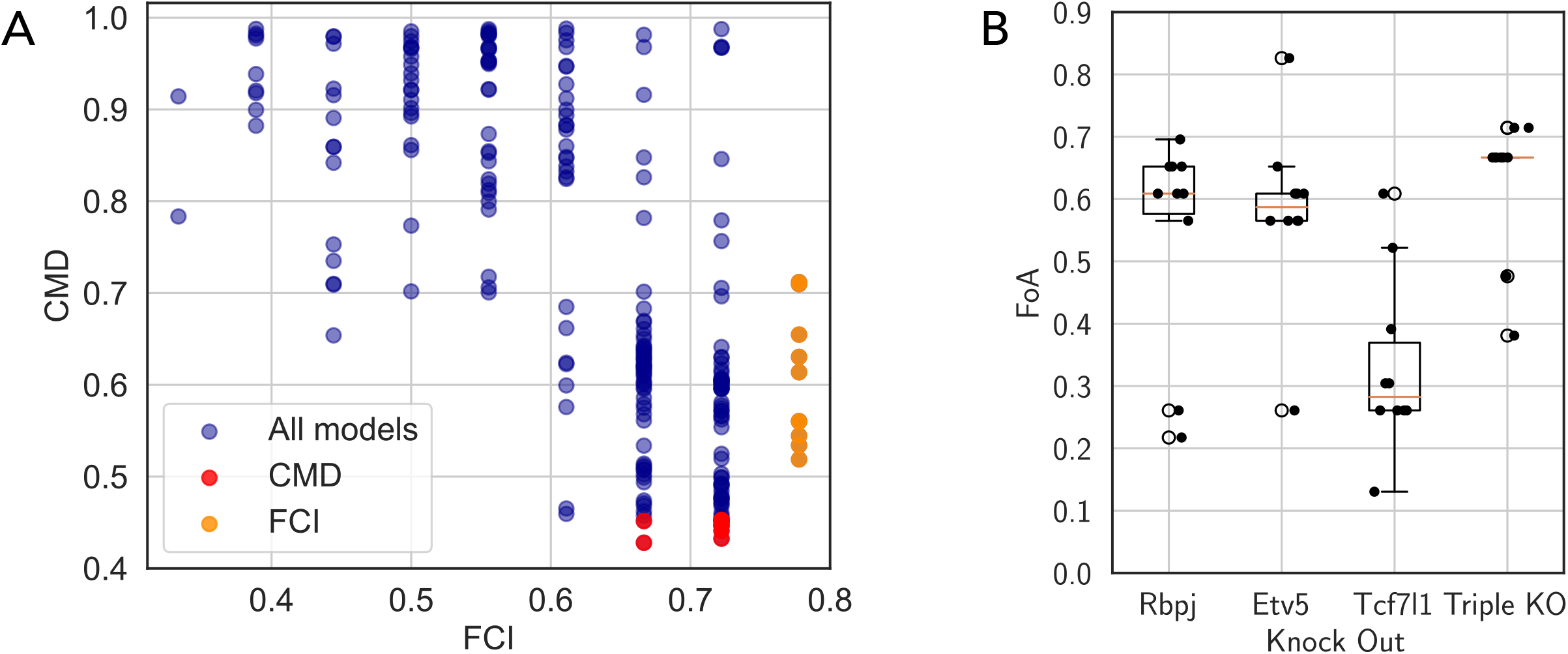
A.Scatter plot of fraction of correctly inferred interactions, FCI, and Correlation Matrices Distance, CMD, for IGNITE-inferred GRNs. Each point represents one of the 250 GRNs inferred using the input dataset (scRNA-seq data with LogNorm, PST, and MB). The 10 models with the lowest CMD are highlighted in red, while the 10 with the highest FCI are marked in yellow. B. Whisker plot of the Fraction of Agreement, FoA, values for each type of KO (Rbpj, Etv5, Tcf7l1 and all three combined) for 10 GRN models inferred with IGNITE that exhibited the highest FCI values (selected from 250 tested models). The box is drawn to represent the lower and upper quartiles, with an orange line indicating the median. The whiskers extend from the box to illustrate the full range of the non-outlier data. Any data points beyond the whiskers are referred to as outliers.

## 4 Discussion

In silico approaches can be applied to modelling and predicting the effects of perturbations on cells, such as knockouts, forced gene activation, or external stimuli. Successful prediction of such effects would allow to significantly scale up the number of inspectable perturbations on a system without the need to perform them in experiments. Hence, an efficient algorithm capable of predicting the effect of perturbations under unseen conditions has the potential to decrease experimental cost, e.g., in drug screening, or to enable customised treatments for individual patients by facing the presence of limited data. Many methods focused on GRN inference [2] or on predicting perturbation effects [11, 1]. However, previous perturbation methods required model training on perturbed data, limiting their scope [1]. Furthermore, a variety of algorithms rely on different types of data as input, downplaying their applicability depending on data availability and compatibility. For instance, algorithms such as RE:IN [3] require ChIP-Seq and bulk transcriptomics data, and need knockout experiments to build the GRN. Understanding how to link cell phenotypes with inferred GRN and how different conditions affect cell behaviour remains challenging.

To address this fundamental biological question, we developed IGNITE, a method that exploits GRN inference to simulate gene perturbations. The significant advantage of IGNITE is its ability to inspect the role of gene interactions in shaping cell phenotypes. During the initial processing stage, IGNITE exploits the temporal features of the scRNA-seq dataset computing the pseudotime. Then, combining it with mini-bulks, it tackles the challenge of high false negative dropout rates by averaging across neighbouring cells along the pseudotime path. For the inference step, IGNITE reconstructs a kinetic Ising model, inferring a signed and directional GRN. This novel technique offers several advantages. (1) It is capable of generating wild-type data, replicating the gene activation patterns within each cell, and (2) simulating GRN perturbations, such as gene knockouts, as well as accurately forecasting their results predicting how KOs alter gene activity within the GRN. These crucial features pave the way for a deeper understanding of the underlying biological mechanisms governing gene interactions. (3) IGNITE starts from unperturbed single-cell RNA sequencing data and thus does not need additional inputs (e.g. ATAC-seq data). (4) With an unsupervised machine learning approach, it infers directed, weighted, and signed GRN that best describes observed cell phenotypes, where patterns of activation among the naive, formative early, formative late, and other gene groups are discernible. We verified that the patterns of generated behaviour are consistent with the experimental predictions for triple KO. Although we did not explicitly carry out simulations of forced activations of a gene, IGNITE can implement this perturbation as well. We highlight that the GRN inferred by IGNITE is *effective*, as it does not describe biological interactions among genes in real cells. Rather, it captures the functional effective organisation in GRN linked to specific cell behaviours. The application of IGNITE to differentiating PSCs underscored its ability to infer effective GRN that successfully forecast cell phenotypes under wild-type or perturbation conditions (single and triple knockouts of Rbpj, Etv5, and Tcf7l1), elucidating the behaviour of PSC during differentiation.

To benchmark IGNITE, we adapted and tested other algorithms to produce comparable outputs. For our differentiation data, MaxEnt [9, 8] was not sufficiently informative as it can not infer the directionality of the network. SCODE [10], designed for GRN inference from scRNA-seq data of differentiating cells, does not model the intrinsic variability seen in the data, as it performs inference with a deterministic model. Furthermore, SCODE does not exploit the model’s ability to generate simulations of system perturbations, even when feasible. We explored this untested SCODE generation capability to simulate single and triple KOs. CellOracle [11] uses GRNs inferred from single-cell multi-omics data to perform in silico perturbations of transcription factors, simulating the resulting changes in cell identity based on unperturbed wild-type data. However, this approach does not describe the underlying process, as it focuses on the propagation of the perturbations in the GRN. We did not employ CellOracle to dissect cell identity via network inference. Rather, we limited its application to measure the effects of KO perturbations, in a way comparable to IGNITE. We included the triple KO simulation, a scenario that was not investigated in the original work. Even though all methods were able to correctly infer known interactions in the GRN, IGNITE outperformed all of them in generating both wild-type and perturbed data. This is because IGNITE leverages its generative non-linear dynamics to learn the properties of the underlying effective GRN, rather than relying on data-constrained GRN inference or perturbation rules to propagate gene expression changes after KOs.

Our results show that IGNITE can provide useful information by modelling GRNs to describe differentiation. Importantly, however, there are limitations to its scope of applicability. First, IGNITE requires a correct pseudotime ordering. This is a crucial step shared by other inference algorithms, such as SCODE, as temporal patterns of gene expression remained consistent after PST ordering. PST and MB allow us to avoid alternative methods based on assumptions that might not precisely capture the actual biological variability, such as the imputation of zero gene expression values implemented by CellOracle. Another important feature of IGNITE is that the underlying Ising model simplifies gene expression information into a dichotomous variable by approximating genes as spins. Although this representation does not appear to limit GRN inference capabilities or prediction of perturbation effects, it is possible to generalise the approach by adopting a model in which variables have multistate values, such as the Potts model [41]. Moreover, several extensions of IGNITE are possible based on relaxing three of the model assumptions: (i) the generated data only describe stationary trajectories by means of a non-equilibrium steady state [12], (ii) the network is static, and (iii) the interactions are only pairwise. We stress that our results show that, even with these assumptions, IGNITE is able to replicate all observed gene activity patterns in the initial data. Furthermore, even if we tested it on a GRN with 24 genes known to be influential in the differentiation process, IGNITE is scalable to larger numbers of genes. In this direction, future developments will have to tame the possibility of multiple local maxima of the likelihood that IGNITE aims to maximise. Our results are promising, since we have found IGNITE to be stable and robust in searching for optimal solutions. This stability is crucial because it implies that IGNITE is capable of distinguishing between truly informative GRNs with biological meaning.

Finally, IGNITE uses scRNA-seq data, which may not allow the inference of specific types of gene interactions. For instance, the effects of a Tcf7l1 KO could not be precisely predicted. This outcome was expected since TCF7L1 is regulated at the level of protein stability and DNA affinity. These aspects cannot be captured by scRNA-seq. The inclusion of other prior knowledge into the model is possible, although we have shown that it does not appear to enhance its performance. This suggests that adding predefined constraints may not always improve performance and could potentially lead to suboptimal solutions. Indeed, the model only provides an effective description of the GRN, and integrating existing knowledge may not always be straightforward or advantageous. This highlights the importance of a cautious approach when integrating known information into such algorithms.

Overall, IGNITE is a valuable machine learning approach for studying PSC differentiation. Its applicability can be generalised to characterise different cell systems and their GRNs without requiring experimental perturbations, as several other models [42]. On the contrary, not only IGNITE relies on unperturbed data, but it also enables reliable in-silico predictions on system perturbations that can be implemented even for genes with an unknown binding motif.By inferring regulatory interactions even for genes with low expression levels, it also addresses the issue of data sparsity inherent in scRNA-seq datasets. This makes IGNITE a powerful tool for navigating the complexities of gene regulatory networks, even in the context of sparse or low-expression data. Furthermore, IGNITE allows exploring how gene interactions shape cell patterns. This is instrumental in gaining a comprehensive understanding of the system and modelling its perturbations solely based on scRNA-seq data, without the need to integrate additional data types.

## 5 Methods

### 5.1 Experimental Procedure

#### 5.1.1 Maintenance of mESCs

Mouse ESCs (R1) were cultured as previously described [43]. Briefly, the base medium is HyClone DMEM/F12 base medium without Hepes (Cytiva), with 4mg/mL AlbuMAX™ II Lipid-Rich Bovine Serum Albumin (GIBCO™), 1× MACS NeuroBrew-21 with Vitamin A (Miltenyi Biotec), 1× MEM NEAA (GIBCO™), 50U/mL Penicillin–Streptomycin (GIBCO™), 1mM Sodium Pyruvate (GIBCO™), and 1×2-Mercaptoethanol (GIBCO™). The base medium was supplemented with 3.3mM CHIR-99021 (Selleckchem), 0.8mM PD0325901 (Selleckchem), and 10ng/mL hLIF (provided by the VBCF Protein Technologies Facility) to create the 2iL self-renewal medium. Cells were cultured on Greiner Bio-One CELLSTAR Polystyrene 6-well Cell Culture Multiwell Plates, pre-coated with Poly-L-ornithine hydrobromide (6 µg/mL in 1xPBS, Sigma-Aldrich, for 1 hour at 37°C, SigmaAldrich) followed by Laminin from Engelbreth-Holm-Swarm murine sarcoma basement membrane (1.2 mg/mL in 1xPBS for 1 hour at 37°C, SigmaAldrich). Routine cells passaging was carried out at a 1:6 ratio every 2 days using 1× Trypsin–EDTA solution (Sigma) at 37°C, and the reaction was stopped by using 10% fetal bovine serum (FBS, Sigma) in the base medium.

#### 5.1.2 10X Genomics scRNA-seq with MULTI-Seq Barcoding and Validation

10X Genomics scRNA-seq was performed as described in [43]. Cells were harvested via trypsinization for 10 mins at 37°C. Trypsinization was stopped using 10% FBS in base medium. Subsequently, the cell pellets were resuspended in PBS to remove any remaining FCS, and their quantification was carried out using a CASY cell counter (Biovendis). Cell labelling with MULTI-Seq barcodes followed a protocol as described in [44]: 0.5 million cells per sample were resuspended in PBS and incubated with a mixture of barcodes and lipid anchors for 5 minutes on ice. Co-anchors were subsequently introduced for an additional 5 minutes, followed by quenching the reaction with 1% BSA/PBS. Cell washing was performed twice using 1% BSA/PBS, and then cells were resuspended in 0.04% BSA. After combining all samples, filtration was carried out through FACS strainer cap tubes and the concentration was adjusted to 1 million cells/mL. Following the manufacturer’s instructions, only pools exhibiting > 80% cell viability were used for the subsequent preparation of the 10X library. For MULTI-Seq identification, libraries were prepared in parallel with cDNA libraries sequenced on Illumina NextSeq550 or NovaSeq platforms. To confirm the specificity of MULTI-Seq, ATTO-488 and ATTO-590 conjugated barcodes were used, as detailed in the [44]. Cell labelling with barcodes followed the aforementioned procedure, with label validation performed through FACS analysis.

### 5.2 scRNA-seq data processing and analysis

#### 5.2.1 The dataset

cDNA reads were mapped to the mm10 reference genome and demultiplexed into droplets by using Cellranger as described in [43]. MULTISeq barcodes were mapped to droplets and counted using CITE-Seq-Count and the Seurat vignette “Demultiplexing with Hashtag oligos” was used to perform cell classification. The cells in the dataset belong to 5 time points (0h, 6, h, 12h, 24h, 48h).

#### 5.2.2 Quality control and cell filtering

To ensure data quality, we removed doublet and negative cells from the analysis. Per-cell QC metrics were computed using the `perCellQCMetrics` function from the `scater` R package. Outlier cells were identified and removed using the `quickPerCellQC` R function with a threshold of 3 median absolute deviations (MAD) from the median. In addition, genes with low expression levels (observed in fewer than 20 cells) were excluded from the analysis. After implementing these steps, the dataset comprised *N*_genes_ = 13833 genes and *N*_cells_ = 9894 individual cells.

#### 5.2.3 Normalisation

We normalised the raw gene expression counts by dividing the total counts for each cell by the corresponding size factor (the sum of the counts for that cell). Subsequently, we log-transformed the normalised expression values by adding *PseudoCount* = 1 and performing a log2-transformation. Therefore, indicating the raw gene expression for the gene *i* in the cell *α* as 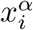 we get its normalised gene expression 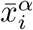 as follows:

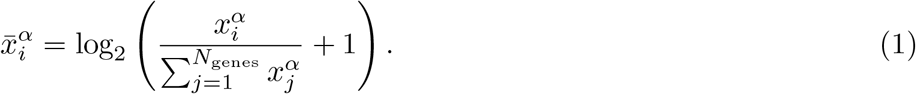

#### 5.2.4 Dimensionality reduction

To transform high-dimensional scRNA-seq data into lower-dimensional representation, we employed t-SNE [31] and UMAP [32] techniques. Initially, for t-SNE dimensionality reduction, we selected a subset of 2318 genes that have been previously established as relevant in the studied differentiation process [6]. Using this gene set, we generated a two-dimensional t-SNE dataset. Then, we projected the t-SNE-transformed data into a 2-dimensional UMAP space (Fig. S1B). The UMAP analysis revealed a coherent temporal progression of cells from the cells sample at *t* = 0 *h* to the sample at *t* = 48 *h* along the first dimension of UMAP. This progression was discernible, with distinct segregation of cells from different time points, forming an ordered sequence from left to right. This observation was further supported by examining the gene expression patterns of our 24 selected genes within the UMAP space (e.g. the three genes in Fig. 1E).

#### 5.2.5 Clustering

We performed a clustering analysis using a graph-based approach on UMAP-transformed data. Initially, we constructed a shared nearest neighbour graph with a total of 30 nearest neighbours (k = 30). We used the `buildSNNGraph` function of the R package `scran`. Next, we applied the Walktrap algorithm to compute the clusters, implemented in the igraph R package. We merged smaller clusters to form larger ones, drawing on time sample labels to understand clusters composed of cells that were generated at the same time point. This procedure was carried out to reinforce both the robustness and interpretability of our analysis on the evolution of cell subpopulations. To validate the biological significance of the resulting clusters, we investigated the distribution of gene expression of the 24 relevant genes within each cluster (e.g. three genes in Fig. S1D). Our exploration uncovered distinct patterns: naïve genes exhibited heightened activity mainly in the left clusters (clusters labelled as 2 and 3), while formative genes demonstrated increased activity predominantly in the clusters in the right part of the UMAP space (clusters 4, 5, and 6).

#### 5.2.6 2Cell-Like Cells cluster

We investigated the cluster 1, as for the 198 cells in this cluster, the gene expression of the 24 selected genes did not allow us to understand whether it was a cluster with a prevalence of naive, formative early or formative late genes. We computed the top 100 most up-regulated genes and the top 100 most down-regulated genes in this cluster concerning all other clusters by using the scran R package function `findMarkers`. Remarkably, we observed that the up-regulated gene set contained genes associated with the 2Cell-Like stage [33]. This finding allowed us to confidently assign an identity to the cells within this cluster as 2CLCs. From this point on, we excluded this cluster from our studied dataset since we are not interested in this cell state.

#### 5.2.7 Pseudotime

We employed the Slingshot algorithm [23] to compute the pseudotime (PST) of cells. This algorithm starts from a low-dimensional space and needs cluster labels for each cell. For this purpose, we utilised the UMAP space data, as well as the clustering labels. We set the input of the slingshot function: we indicated cluster “2” as the start cluster and cluster “6” as the end cluster. This choice was made after considering the high gene expression regions in the UMAP space for each of the 24 selected genes. Moreover, we observed that the real temporal progression of the process described by the sampling times and the PST progression are in agreement by generating the violin plots of the pseudotime values for the cell in each time point separately (Fig. S1E).

#### 5.2.8 Gene selection

The dataset after the pseudotime implementation includes 13833 genes and 9696 cells. We focused only on 24 genes known from the literature as relevant and informative in the naive to formative transition of PSCs [3, 6, 29, 30].

#### 5.2.9 Mini-Bulk

We called Mini-Bulk (MB) the approach we chose to compute the moving average over the cells in the dataset: we averaged subsets of cells with a window size of 150 cells and a step size of one. The choice of this value for the step size ensures that the dimensions of the resulting dataset are comparable to those of the original one. We applied the MB procedure to our single-cell dataset, normalised and with cells ordered by following the pseudotime values. In the end, we obtained a dataset of 24 genes and 9547 cells (Fig. S1 F). The dataset thus obtained will be the input for all GRN construction methods, except for CellOracle, which does not require PST and MB

### 5.3 Bulk RNA-seq data processing

We utilised RNA sequencing data derived from embryonic stem cell (ESC) experiments as described by Carbognin et al. [6]. To process the data, we followed the same procedures detailed in the original study. The initial step involved converting the Illumina NovaSeq base call files to fastq files using the bcl2fastq tool (v2.20.0.422). The raw data were then analysed using the Next Generation Diagnostic srl proprietary 3^*’*^DGE mRNA-seq pipeline (v2.0). This pipeline includes quality filtering and trimming steps using the bbmap suite (v37.31) from the Joint Genome Institute (JGI). The aligned reads were mapped to the mm10 reference genome assembly (release 76) and counted by gene using the mm10 Ensembl assembly (release 93). We applied the Counts per Million (CPM) normalisation approach to the RNA-seq datasets. Then we averaged the two samples produced under the same conditions at the same time step.

### 5.4 IGNITE

We assumed that the activity of the genes can be effectively captured by a kinetic Ising model, governed by a simple spin-flip dynamics known as Glauber dynamics [12]. To infer the GRN interactions, we assessed the inverse problem for this model. Following the work of Zeng et al. [13], we further assume that the spins are updated asynchronously, and that the whole GRN is a fully connected network, with 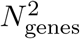 possible interactions.

#### 5.4.1 Gene activity: Binarisation of the inputs

We represent genes as a set of spins 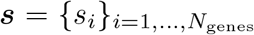 where each *s*_i_ is a binary variable that can assume values +1 and −1. Therefore, in order to infer the GRN from an Ising model and run IGNITE, it is necessary first to binarise transcriptomic data. We started from the input dataset (log-normalised scRNA-seq matrix with PST and MB). If we denote with *x*_*i*_(*t*) the gene expression of gene *i* for the cell with PST value *t*, the binarisation of *x*_*i*_(*t*) was performed as follows:

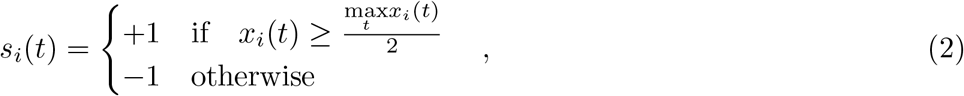

where *s*_*i*_(t) = +1 if gene *i* is active, and *s*_*i*_(t) = −1 if it is inactive. Based on gene expression, this approach computes gene activity (GA) as a binary measure of activity, assigning a value of −1 for inactive genes and +1 for active genes. We tested this approach by verifying that naive genes are mainly active at low PST values, whereas formative genes are mainly active during later steps in the pseudotime temporal evolution computing the heatmap of GA (Fig. S1G).

#### 5.4.2 Asymmetric kinetic Ising model

The kinetic Ising model is specified by a coupling matrix of entries *J*_*ij*_, which quantify gene-to-gene interactions and describes the GRN, and by a set of external fields *h* = {*h*_*i*_}, which measure the bias for a given gene to be active (*h*_*i*_ > 0) or inactive (*h*_*i*_ < 0). From the timeseries of *s*_*i*_(*t*), the goal of IGNITE is to infer *J*_*ij*_ and *h*_*i*_, which are assumed to be constant in time. Both *h*_*i*_ and *J*_*ij*_ are expressed in units of temperature.

Consider a time discretization with steps of size *δt*. During the time evolution of the model, the spins *s*_*i*_ are updated asynchronously. At each time step *t*, a spin *i* is randomly selected with probability *γ*_*i*_*δt*, and its value *s*_*i*_ is updated according to the transition probability:

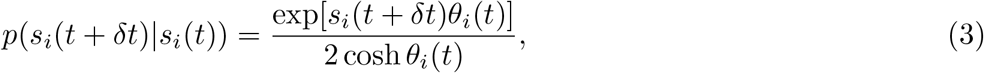

where *p*(*s*_*i*_(*t* + δ*t*)|*s*_*i*_(*t*)) is the probability of the new spin value at time *t* + δ*t* given the one at the previous time *t*, and *θ*_*i*_(*t*) = *h*_*i*_ *+∑*_*j≠i*_ *J*_*ij*_*s*_*j*_(*t*) is the effective local field acting on the *i*-th spin at time *t*.

#### 5.4.3 Model reconstruction from data

In principle, to perform the inference, we would need to know both the spin values at each time, s = {s_*i*_(*t*)}, and the update times, ***τ*** = {***τ*** _*i*_}, for a given time interval *T* with timestep δ*t*. Then, starting from the spin history, our goal is to reconstruct the model parameters *h*_*i*_ and *J*_*ij*_. We can determine the learning rules for these parameters by likelihood maximisation [13]. We can write the likelihood as:

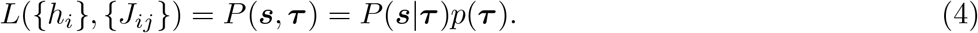

For each spin, the update times are a discretised Poisson process. This means that every time step *t* is an element of the update time set ***τ*** with probability *γ*_*i*_δ*t*. Therefore, the update time probability *p*(***τ***) is independent of the model parameters that we want to infer. For simplicity, we take *γ*_*i*_ = *γ*, assuming that *γ* is known a priori, so that we do not need to determine it during the inference problem. For simplicity, we set *γ* = 1. Hence, the likelihood maximisation can be carried out over *P*(***s***|***τ***) only.

Marginalising out the update times, we can obtain a simpler form of the update probability for the *i*-th spin, which is exactly the Glauber dynamics update rule:

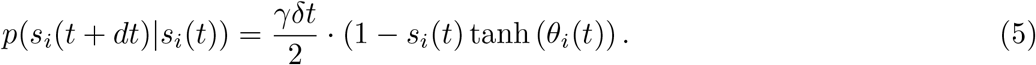

Then, the logarithm of the likelihood, i.e. the log-likelihood, is:

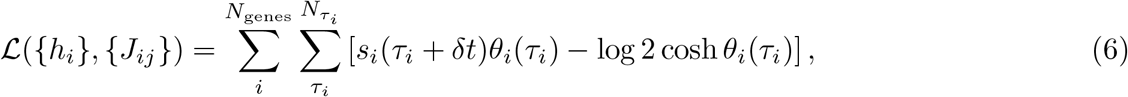

where 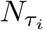 is the number of update times for the gene *i*. We stress that maximising the log-likelihood is equivalent to maximising the likelihood. By taking a partial derivative of Eq. 6, we obtain the following learning rule for the interaction matrix:

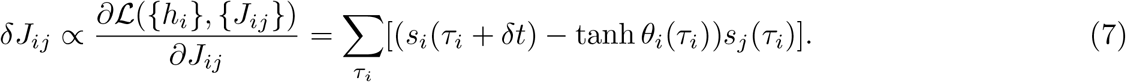

Importantly, if we define *J*_*i0*_ = *h*_*i*_ and *s*_0_(*t*) = 1, Eq. 7 includes also the learning rule for fields *h*_*i*_ [13].

Crucially, Eq. (7) requires the knowledge of the update times for the spins, which we typically cannot access. Indeed, in each time interval, one spin is randomly chosen for updating, but it does not necessarily flip. Thus, it is not generally possible to measure the update times from the timeseries only. To overcome this problem, we can proceed as in the work by Zeng et al. [13]. We define the correlation function C_*ij*_(*t*) *≡* ⟨*s*_*i*_(*t*_0_ +*t*)*s*_*j*_(*t*_0_) ⟩, where ⟨·⟩ is the average over all realisations of the stochastic dynamics. The time derivative of the correlation function is:

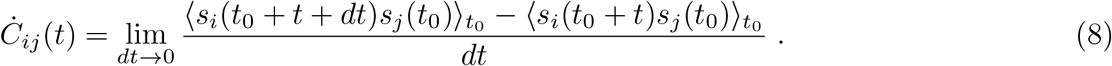

By noting that we can separate the terms with and without spin flips, we can write:

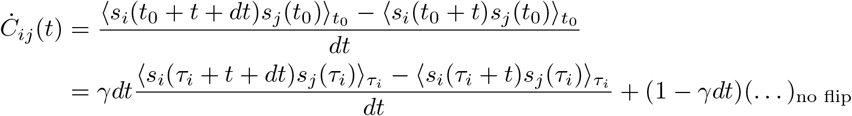

The term without flip vanishes because the spin did not update, and therefore *s*_*i*_(*t* + *dt*) = *s*_*i*_(*t*). Hence, since δ*t* is the smallest time interval available from experimental trajectories, we can write:

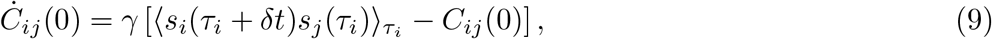

so that

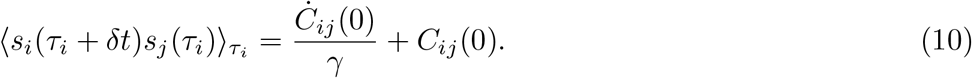

The updating rule (Eq. 7) can be rewritten by substituting the first term of the right part of the equation with Eq. 10:

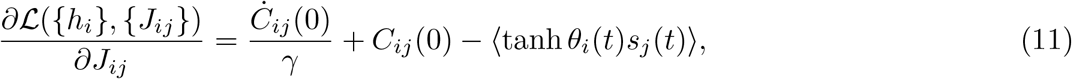

where the second term in the right part of Eq. 11 is an average over all time points. We note that this term is equivalent to the average over *τ*_*i*_ in Eq. 7, since tanh *θ*_*i*_(*t*)*s*_*j*_(*t*) is insensitive to whether an update has been made. Since the choice of the unit of time intervals is not relevant for inference, we set *δt* = 1 without loss of generality. Although convergence to the global maximum cannot be guaranteed, the update rule in Eq. (11) will reach a local maximum that corresponds to a coupling matrix representing a functional GRN.

We note that the interaction matrix *J* includes self-couplings for all genes. However, measuring mRNA does not allow us to distinguish between target and regulator genes. Thus, self-couplings lack direct biological relevance. However, this computational inclusion, while not reflecting direct biological significance, stabilises the inference process by introducing an informative bias that could compensate for specific dynamics or interactions not directly captured by the model. For this reason, we infer the self-couplings, and then remove them from the GRN without further investigation.

#### 5.4.4. Optimisation algorithms

To find the maximum of Eq. 11, we implemented two optimisation methods, the Momentum Gradient Ascent (MGA) algorithm (implemented as the maximisation version of the Stochastic Gradient Descent algorithm [45]) and the Nesterov-accelerated adaptive moment estimation algorithm (NADAM) [46]. In general, the maximisation of the likelihood is performed starting from an estimate of *J*_*ij*_ and *h*_*i*_, and updating this estimate with a small value in the direction of the gradient. This is repeated until convergence. Each loop of the learning rule is called an epoch.

##### Momentum Gradient Ascent algorithm

For the MGA algorithm, we adopted a step-decaying learning rate and an *L*_1_ regularisation. The learning rate for the *n*-th epoch is:

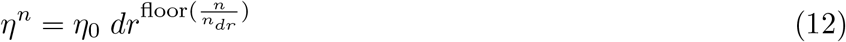

with *η*_0_ initial learning rate, and *dr* drop of the learning rate every *n*_*dr*_ epochs. The *n*-th optimisation step for the fields can be written as:

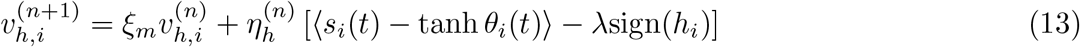

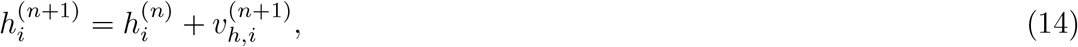

and the *n*-th optimisation step for the couplings is:

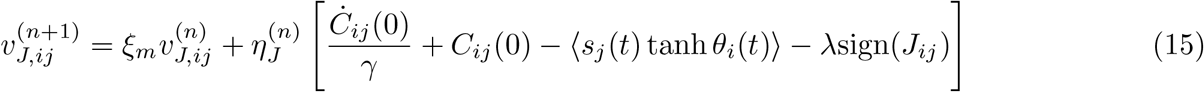

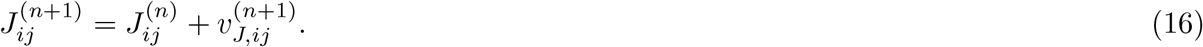

The parameter *ξ*_*m*_ in Eq. 13 and Eq. 15 is the momentum parameter, with 0 *≤ ξ*_*m*_ *≤* 1, and *λ* is the regularisation parameter.

##### Nesterov-accelerated Adaptive Moment Estimation algorithm

With the NADAM algorithm, we implemented both a step-decaying learning rate (Eq. 12), and *L*_2_ regularisation. To compute the first-moment m and the second-moment *v* of the gradient with respect to the coupling and the fields, we can write the *n*-th optimisation step as follows:

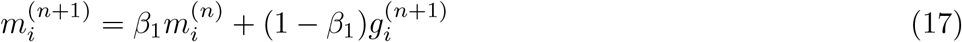

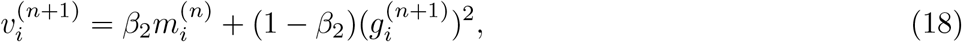

where *g*_*i*_ is either 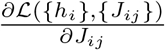 or 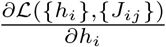 (see Eq. 11). To perform the learning step, we need to introduce the bias-corrected moments:

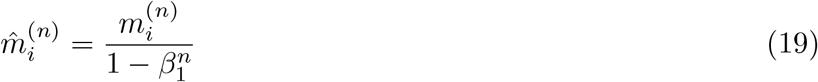

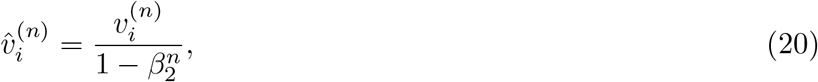

so that:

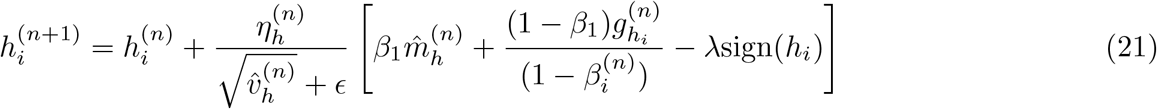

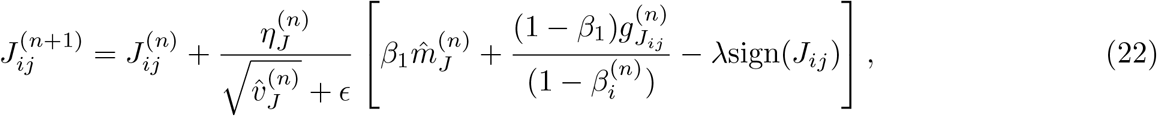

where *λ* is the *L*2 regularisation parameter, and, *β*_1_ = 0.9,, *β*_2_ = 0.999 and ϵ = 10^*−*8^ are fixed parameters of the NADAM algorithm.

#### 5.4.5 Hyperparameters of the optimisation algorithms

To perform the GRN reconstruction from Eq. 11 we have to set the following hyperparameters in both the MGA and the NADAM optimisation algorithms.

- For the MGA optimizer: the step-decaying learning rate parameters, *η, dr* and *n*_*dr*_, the momentum parameter *ξ*_*m*_, and the *L*1 penalty term *λ*.
- For the NADAM optimiser: the step-decaying learning rate parameters, *η, dr* and *n*_*dr*_, and the *L*2 penalty term *λ*.

We also note that the optimisation problem is not convex, as we expect Eq. 6 to be characterised by many local maxima. To avoid sub-optimal solutions, we performed a random search among *N*_trial_ = 250 randomly selected sets of hyperparameters. The possible values considered for the hyperparameters are shown in the Tab. 4. The best set of hyperparameters can be defined with or without using prior knowledge.

**Table 4:**
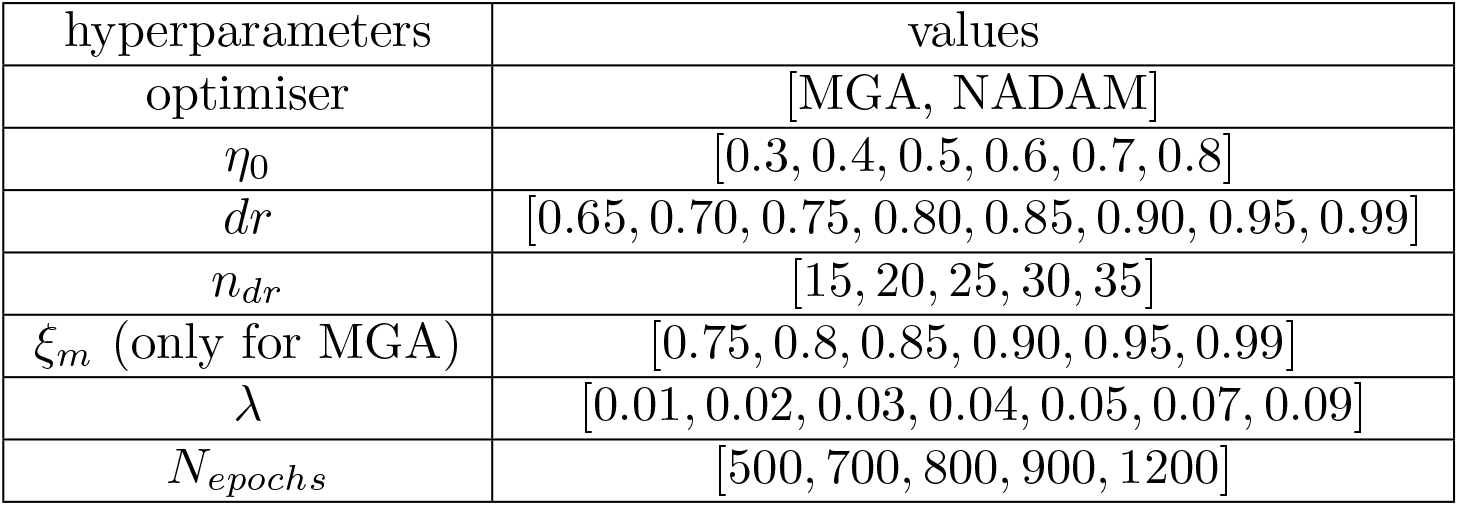
Set of hyperparameters values for the random search. The momentum parameter is used only if the MGA optimiser is selected.

i. With prior knowledge: we know a set of 18 interactions and we select the set of hyperparameters for which the measured FCI is maximum. If there are models that have equal FCI, we randomly select one of these models.
ii. Without prior knowledge: we select the set of hyperparameters for which the measured CMD is minimum. If there are models that have equal CMD, we select randomly one of these models.

#### 5.4.6 Comparison with null models

We conducted a permutation test to assess the statistical significance of the inferred interactions. The permutation test involved shuffling the row and column indices of the gene expression data *N*_test_ = 50 times and then evaluating *N*_sets_ = 50 sets of hyperparameters for each test dataset, to ensure thorough testing of different hyperparameter configurations. The final number of inferred null GRN with IGNITE is 2500. We investigated the statistical significance of the GRN inferred using IGNITE. We hypothesised that a network derived by chance would exhibit interaction values similar to those of null models. We assessed the 12 known interactions correctly inferred with IGNITE by comparing their values from the actual GRN against the probability distributions of these null model GRNs. Specifically, we checked if the values predicted by IGNITE for these interactions lay outside the 5th to 95th percentile range of the shuffled data. Since only three interactions aligned with the null model expectations (Nanog-Otx2, Stat3-Gbx2 and Tcf7l1-Esrrb), we reject the null hypothesis, supporting the statistical significance of the IGNITE GRN.

### 5.5 MaxEnt reconstruction

The element *ρ*_*ij*_ between the genes *i* and *j* of the MaxEnt interaction matrix can be calculated as:

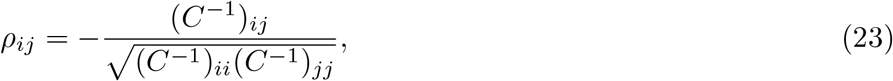

where C^*−*1^ is the inverse matrix of the covariance matrix calculated starting from our transcriptomics datasets, as described in [9, 8]. The interaction matrix *ρ* is symmetric.

### 5.6 SCODE reconstruction

SCODE [10] is a method for inferring regulatory networks from scRNA-Seq data designed for differentiating PSCs. It utilises linear ordinary differential equations (ODEs) and linear regression to capture the observed gene expression dynamics. We selected this method as the gold standard method comparable to IGNITE, because, as highlighted in [2], it has characteristics similar to those of IGNITE: (i) it needs time-ordered cells to capture the network dynamics, (ii) the inferred interactions are directed (asymmetric interaction matrix) and (iii) signed, and (iv) it is possible to generate new data in wild-type or under perturbation conditions. Looking at the other models evaluated in [2], SCODE is the model with these characteristics and with the best accuracy results for the experimental scRNA-seq datasets.

SCODE efficiency is based on the fundamental concept that the patterns of expression dynamics are finite and that expression dynamics can be accurately reconstructed using a limited number of these patterns. Therefore, the expression dynamic vector should be reduced from a length of *N*_genes_ genes to a length *D*, with *D ≪N*_genes_. This number *D* is a parameter of the model and its value should be optimised, as suggested by the authors. To do that, we divided the dataset in training (*N*_genes_ × *N*_cells_ = 24 × 7638) and test datasets (*N*_genes_ × *N*_cells_ = 24 × 1909, 20% of the total dataset cells). We applied SCODE to training data and we evaluated the validity of the optimised model by computing the residual sum of squares (RSS) of the test dataset for various values of D (*D* = 2, 4, 6 and 8). For each D, as in the original work, we executed SCODE 100 times independently. We observed that the median of the RSS values was almost saturated at *D* = 6. As suggested by the authors, we checked the reproducibility of the inferred interaction matrix. The correlation coefficient was calculated among the optimised interaction matrices for the 50 replicates with the lowest RSS values, using the test data for each D. The correlations among replicates are high till *D* = 6 and therefore an optimised interaction matrix remains stable till *D* = 6. Considering the saturation of RSS values for the test data and the stability of the estimated interaction matrices at *D* = 6, we selected this value for our analyses. The final inferred GRN interaction matrix is the element-by-element mean of the interaction matrices of the top 50 replicates.

### 5.7 CellOracle application

CellOracle [11] is a machine-learning framework that leverages GRNs inferred from single-cell multi-omics data to simulate gene perturbations and their effects on cell identity. The in silico perturbation unfolds in four steps: (1) Generation of cell-type or cell-state specific GRNs through cluster-wise regularised linear regression on multi-omics data. (2) Estimation of target gene expression changes due to transcription factor (TF) perturbations, using GRN models to simulate the cascading effects of these changes. CellOracle perturbation modelling is deterministic. (3) Calculation of cell-identity transition probabilities by comparing the gene expression shift with that of neighbouring cells. (4) Transformation of these probabilities into a weighted vector, simplifying the complex gene expression shift into a two-dimensional representation to predict cell-state transitions post-TF perturbation. To compare CellOralce with IGNITE and SCODE, we focused on the first two steps of CellOracle, obtaining as output the inferred GRNs and the simulated gene expression after KO perturbations. CellOracle, as it is designed, can generate only perturbed gene expression data, not WT data. CellOracle requires as input the procesed scRNA-seq dataset with dimensionality reduction and clustering implemented. We took as input the scRNA-seq processed as described in 5.2 till the removal of the 2CLCs cluster. The processed dataset has 9696 cells and 2078 genes. These genes, identified as significant in the differentiation process by Carbognin et al. [6], were chosen over highly variable ones due to their established relevance to the process under study. We used as base GRN a fully connected network, inferring with CellOracle the weights of these connections (excluding self-loops, that are not inferred with CellOracle). We did not use the prebuilt GRN as suggested by the authors because in this network most of the interactions known from the literature are not present (78%). Then, we generated perturbed KO data using the function of the CellOracle Python package `simulate shift`. After GRN inference and data generation, we focused only on 24 genes deemed critical for our study, along with their respective GRN.

### 5.8 Fraction of Correctly Inferred Interactions (FCI)

The Fraction of Correctly Inferred Interactions (FCI) is computed from the inferred interaction values. The FCI is given by:

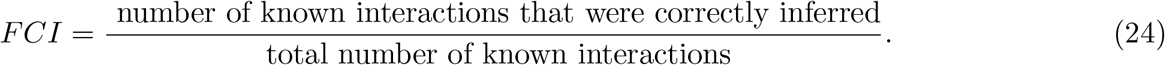

The numerator is determined by counting the instances in which the sign of the inferred interaction values matches the interaction signs known from the literature. The denominator is equal to 18.

### 5.9 Generation and evaluation of WT data

#### 5.9.1 Gene activity generation with IGNITE

We generated gene activity data using the interaction matrix inferred with the IGNITE method. The genes in this dataset, as for the IGNITE inputs, were represented as spins. To generate a new dataset, we leveraged the Ising model parameters, the inferred interaction matrix (*J*) and external fields (*h*), to determine the gene states for different cells by using the Glauber spins update rule. The resulting dataset is a matrix with the structure of the original data, with rows representing genes and columns representing distinct cells corresponding to different time steps. The data dimensions were equal to the input data ones (*N*_genes_ × *N*_cells_ = 24 × 9547).

To generate the data, we initialised the gene expression state vector for each gene in the network with random spin values. At each time step, we updated the values of each spin independently based on its interactions with other spins. Following Glauber dynamics, every spin can flip with probability 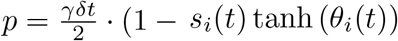 with γ = 1, *δt* = 1, *s*_*i*_(*t*) current spin state of gene *i* at time *t*, and 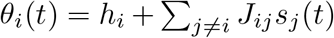 total field acting on spin *i*. Each newly generated dataset is distinct from the others, as IGNITE simulates gene activity via a stochastic dynamics.

#### 5.9.2 Gene expression generation with SCODE

We employed the code provided by the authors of SCODE to reconstruct the expression. We generated a new dataset, capturing the dynamics of the 24 selected genes over 100 time steps, as suggested by the authors. The initial gene expression values for each gene were set to the mean expression of the first 1000 cells from the original log-normalised scRNA-seq dataset with PST and MB. We generated only one dataset since the data generation process is deterministic, ensuring that all generated datasets would be identical.

#### 5.9.3 Correlation matrices distance

We assessed the similarity between the generated WT datasets (using IGNITE or SCODE) and the input dataset as follows.

i. Using IGNITE we generated *N*_trial_ = 250 gene activity datasets with dimensions identical to the original gene activity dataset (*N*_genes_ = 24 and *N*_cells_ = 9547). SCODE is deterministic, therefore we generated only one gene expression dataset with the number of cells suggested by the authors (*N*_genes_ = 24 and *N*_cells_ = 100).
ii. We computed the Pearson correlation matrices for the input datasets and for each of the simulated ones.
iii. For IGNITE only, we computed the element-by-element average of the *N*_trial_ correlation matrices from the simulated data, obtaining an averaged correlation matrix.
iv. We computed the distance between the input data and the generated data matrices as follows:

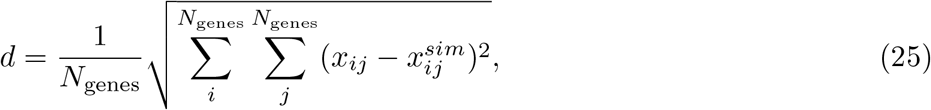

where *x*_*ij*_ represents an element in the correlation matrix of the input gene activity/expression data, and 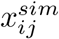 represents an element in the correlation matrix (averaged for IGNITE) of the simulated data.
v. To scale distance measures *d*, we created *N*_shuffle_ = 250 random datasets by shuffling the input gene activity and gene expression datasets, as previously described for the IGNITE null model. We then scaled *d* by dividing it by the average distance between the correlation matrix of the input data and the average correlation matrix of the null model datasets. We call the scaled *d* quantity Correlation Matrices Distance (CMD).

A good inference method will produce simulated data with a CMD smaller than one. A value of CMD = 1 would imply that the simulated data correlation matrices are comparable to what could occur by chance. To quantify the goodness of IGNITE and SCODE results, we performed two statistical tests: (i) a two-sample t-test for IGNITE to compare the distribution of the *d* values of the IGNITE generated data with those of the null model. (ii) the z-score to compare the single *d* values obtained from SCODE with the distribution of the *d* values for the null model.

### 5.10 Clustering of the data

We clustered the cells within a dataset by grouping cells based on similarities in their patterns. This allowed us to explore the underlying structure of the gene expression or gene activity datasets. We started with linkage analysis of the datasets using the `scipy.cluster.hierarchy.linkage` function from the SciPy library. We employed the linkage analysis with the Ward variance minimisation algorithm. This is a hierarchical agglomerative approach aimed at minimising the variance within each cluster. Subsequently, the results of the linkage analysis were visualised using a dendrogram and a heatmap. The heatmap provided an intuitive representation of gene expression/activity patterns across the cellular population. The dendrogram represents the hierarchical structure of the clusters, illustrating how individual cells or groups of cells are merged into larger clusters based on their gene expression/activity similarities (Fig. 2D for input gene activity for IGNITE, Fig. 2E for IGNITE generated gene activity, Fig S2C for input gene expressio for SCODE, and Fig. S2D for SCODE generated gene expression).

### 5.11 Perturbation of the system: gene knockout analysis

We perturbed the system by implementing single gene (for Rbpj, Etv5, or Tcf7l1) and triple gene (for Rbpj, Etv5, and Tcf7l1 simultaneously) knockout simulations to evaluate the prediction capability of IGNITE, SCODE, and CellOracle algorithms.

#### 5.11.1 Generation of KO data

##### With IGNITE

To generate perturbed data after KOs of single or triple genes, we removed the rows and columns of the KO genes from the inferred interaction matrix. Then we generated *N*_trial_ = 250 gene activity datasets as described above (Method section 5.9.1). The datasets have *N*_genes_ = 24 − *N*_KO_ genes, with *N*_KO_ number of KO genes. The number of cells (*N*_cells_ = 9547) is identical to that of the input dataset.

##### With SCODE

We generated the simulated KO by removing the KO gene from the interaction matrix inferred with SCODE. Then we generated one gene expression dataset with *N*_genes_ = 24 *N*_KO_ for each KO, following the procedure described in the Method section 5.9.2. The number of cells is *N*_cells_ = 100, as suggested by SCODE authors. We generated only one dataset since SCODE has deterministic dynamics.

##### With CellOracle

We generated the simulated KO data by using the specific function of the CellOracle Python package `simulate shift`. As for SCODE, for each CellOracle KO, we generated a gene expression dataset with *N*_*genes*_ = 24 − *N*_*KO*_. The number of cells is equal to the input dataset one (*N*_cells_ = 9696). We generated only one dataset since the perturbation procedure for CellOracle is deterministic.

#### 5.11.2 Measuring the influence of the KO gene on the GRN

To assess the impact of gene knockout on gene activity or expression patterns, we compared the WT and KO datasets, considering KOs of single and triple genes. As WT data for IGNITE we generated 250 datasets as detailed in 5.9.1, and for SCODE we generated one WT dataset as described in 5.9.2. For CellOracle the WT dataset is the one used as input to infer the GRN, which is the original processed data with KNN-imputed values for dropout data. To compare the WT datasets to the datasets of the KOs we removed the data of the perturbed genes. The quantification of this influence was carried out as follows.

##### Simulated KO and WT data

We first averaged the gene activity/expression levels per gene across all cells (and in all IGNITE-generated datasets). Next, we calculated the difference between KO and WT gene activity/expressions for each gene *i, Δx*_*i*_:

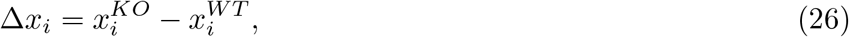

where 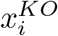 represents the average gene activity/expression of gene *i* in the KO dataset, and 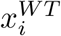 represents the same quantity in the WT dataset. A negative value of Δ*x*_*i*_ indicates that the KO perturbation causes a decrease in the average gene activity/expression of gene *i*, while a positive value suggests that KO causes an increase in the gene activity/expression for this gene. Δ*x*_*i*_ = 0, instead, suggests that KO did not affect gene activity/expression of gene *i*.

##### Experimental data

Given the perturbation of a single gene *j*, or three genes {*j, k, l*}, to measure the effects of the perturbation on experimental data we employed the log2FC measures from [30], with log2FC_*i*_ for gene *i* defined as:

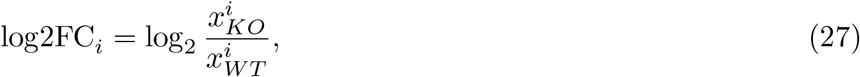

where *x*^*i*^ indicates the gene expression of gene *i*. A negative value signifies a reduction in gene *i* expression due to the perturbation, while a positive value indicates an increase. Log2FC values near zero imply a negligible effect on gene *i*.

For the triple KO experimental data, we used the dataset from [29]. To quantify the perturbation impact on gene expression, we calculated the difference between two log2FC values for each gene: one comparing triple KO to control (2i) conditions, and another comparing gene expression in WT at 72h after the removal of 2i versus 0h. This is still a log2FC measure and improves the effects of the impact of KO on gene expression relative to the baseline expression in the WT condition.

#### 5.11.3 Compare KO measures

To ensure a meaningful comparison between the Δ*x* values of the simulated data and the log2FC values of the experimental data, we normalised both these quantities. For the generated data, all the Δ*x* values obtained from the four KO simulations were divided by the maximum absolute value of Δ*x*, which was calculated across all Δ*x* values from all knockouts. Similarly, both single KO and triple KO Log2FC values were divided by their respective maximum absolute values. This normalisation procedure ensures that all values are comparable and fall within the [−1, 1] interval. Notably, we avoided dividing the Δ*x* and log2FC values by their common absolute maximum value, as the distinct procedures used to generate the two datasets could make direct comparisons inappropriate. Moving forward, we will refer to both normalised values (Δ*x* and log2FC) as scaled KO-WT difference, 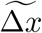 for clarity.

##### Fraction of Agreement (FoA) for KO experiments

To evaluate the concordance between the simulated and experimental KO-WT variation measures, 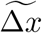 we defined the Fraction of Agreement (FoA). To compute the FoA, we set a threshold value thr = 0.05. Subsequently, we categorised the difference values into three classes: positive if 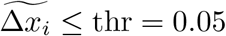 negative if 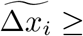 -thr, and null if 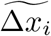 fall within the range (-thr, thr).

Then we counted the instances in which the generated and experimental values of 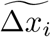 for each gene *i* matched in the categorical range (positive, negative, or null). This count was then divided by the total number of genes to obtain the FoA value (*N*_totalgenes_ = *N*_genes_ N_KOgenes_, equal to 23 for single KO simulations and to 21 for the triple KO).

To evaluate the statistical significance of the observed FoA values against random chance, we used a binomial distribution as a null model. The number of trials is *N*_trials_ = *N*_totalgenes_. As before, we defined as success the instance where the generated and experimental 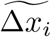 values for a gene fall within the same categorical range, with the potential number of successes varying from 0 to *N*_trials_. The probability of success in any given trial is *p* = 1/3. To determine the significance threshold for the FoA values, we calculated the minimum FoA value needed such that the cumulative probability of observing an FoA value that high or higher, based on the binomial distribution, is less than or equal to 0.05. This threshold defines the value below which a FoA value is considered statistically significant against random chance. The determined threshold is FoA_thr_ = 0.48, for single and triple KO cases.

