## Supplementary figures for "Unveiling Gene Perturbation Effects through Gene Regulatory Networks Inference from single-cell transcriptomic data"

### FIGURE S1

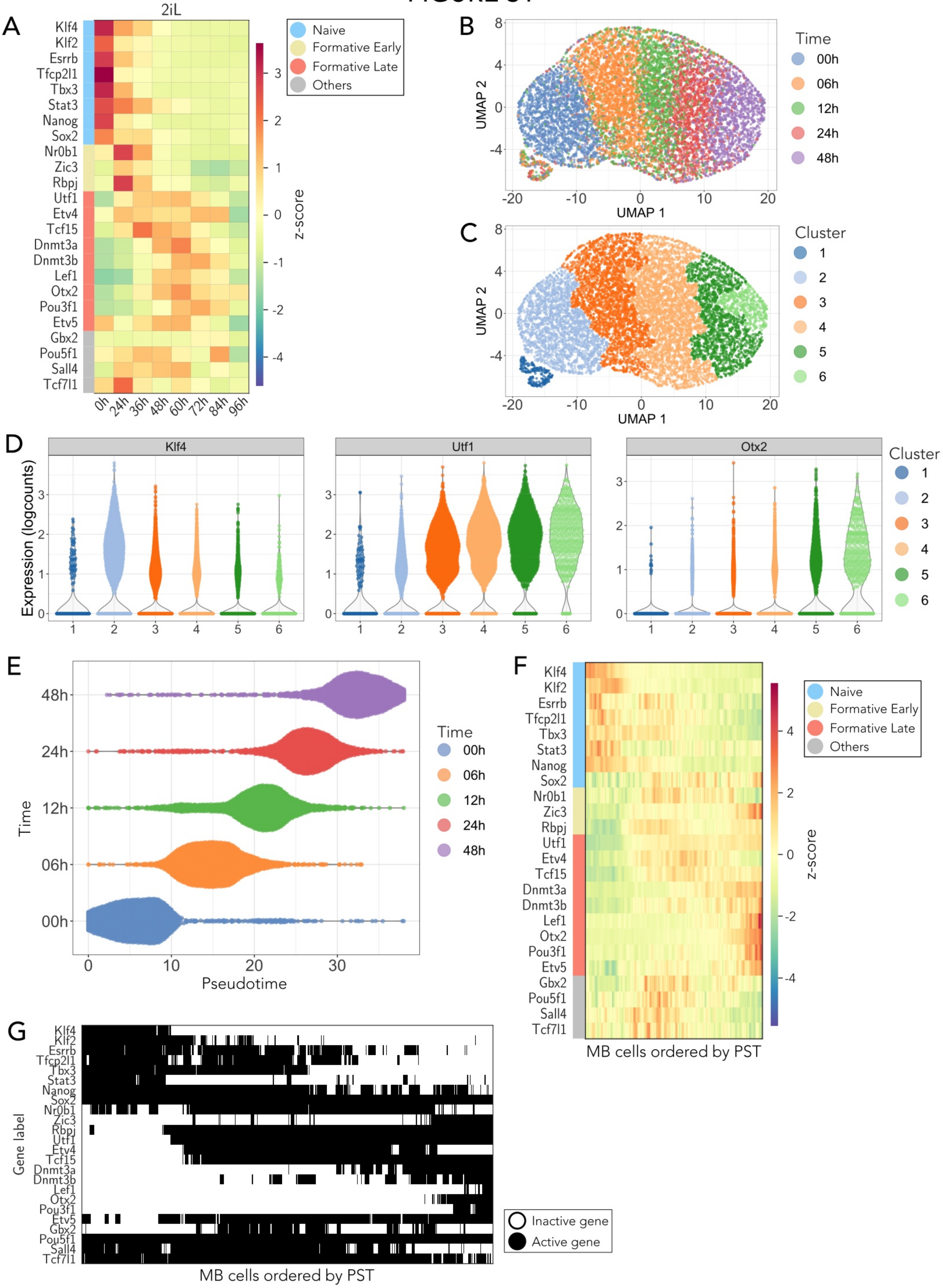
A
2iL
Klf4
Klf2
Esrrb
Tfcp2l1
Tbx3
Stat3
Nanog
Sox2
Nr0b1
Zic3
Rbpj
Utf1
Etv4
Tcf15
Dnmt3a
Dnmt3b
Lef1
Otx2
Pou3f1
Etv5
Gbx2
Pou5f1
Sall4
Tcf7l1
0h 24h 36h 48h 60h 72h 84h 96h
z-score
Naive
Formative Early
Formative Late
Others
B
UMAP 1
UMAP 2
Time
00h
06h
12h
24h
48h
C
Cluster
1
2
3
4
5
6
D
Klf4
Utf1
Otx2
Expression (logcounts)
E
Time
Pseudotime
F
MB cells ordered by PST
G
Gene label
Inactive gene
Active gene
MB cells ordered by PST

#### Figure S1

A. Gene expression z-score of the bulk dataset from [6]. At time 0h the cells are in 2i+LIF (2iL). For each gene, the average is computed over two samples belonging to the same time point. Genes labelling as in Fig. 1C.

B. Visualisation of two-dimensional UMAP of the input experimental dataset (scRNA-seq data with Log-Norm). Each point represents an individual cell coloured according to its sampling time. UMAP1 and UMAP2 indicate the two dimensions of the UMAP space.

C. Two-dimensional UMAP of the input dataset as in Fig. S1B, but with cells coloured according to its custering label. UMAP1 and UMAP2 indicate the two dimensions of the UMAP space.

D. Violin plots of gene expression levels across the six different clusters for three genes (Klf4, Utf1 and Otx2) of the input dataset. Each violin plot illustrates the distribution of log-normalised gene expression values from scRNA-seq data, with the width of the plot is proportional to the density of data points at different expression levels.

E. Violin plot visualisation of pseudotime distributions across different sampling time points for the input experimentaldataset. Each plot corresponds to a sampling time (0h, 6h, 12h, 24h, 48h), with the colours indicating the specific time points. The width of each violin reflects the density of cells at each pseudotime interval.

F. Gene expression z-score of the input dataset (scRNA-seq with LogNorm) with pseudotime (PST) and Mini-Bulk (MB). Genes labelling as in Fig. 1C.

G. Heatmap of the gene activity computed from the input data, where genes are categorised as active (black) or inactive (white) based on a threshold set at half of the maximum expression value for each gene. Rows represent individual genes, while columns represent the cells after PST and MB implementation.

### FIGURE S2

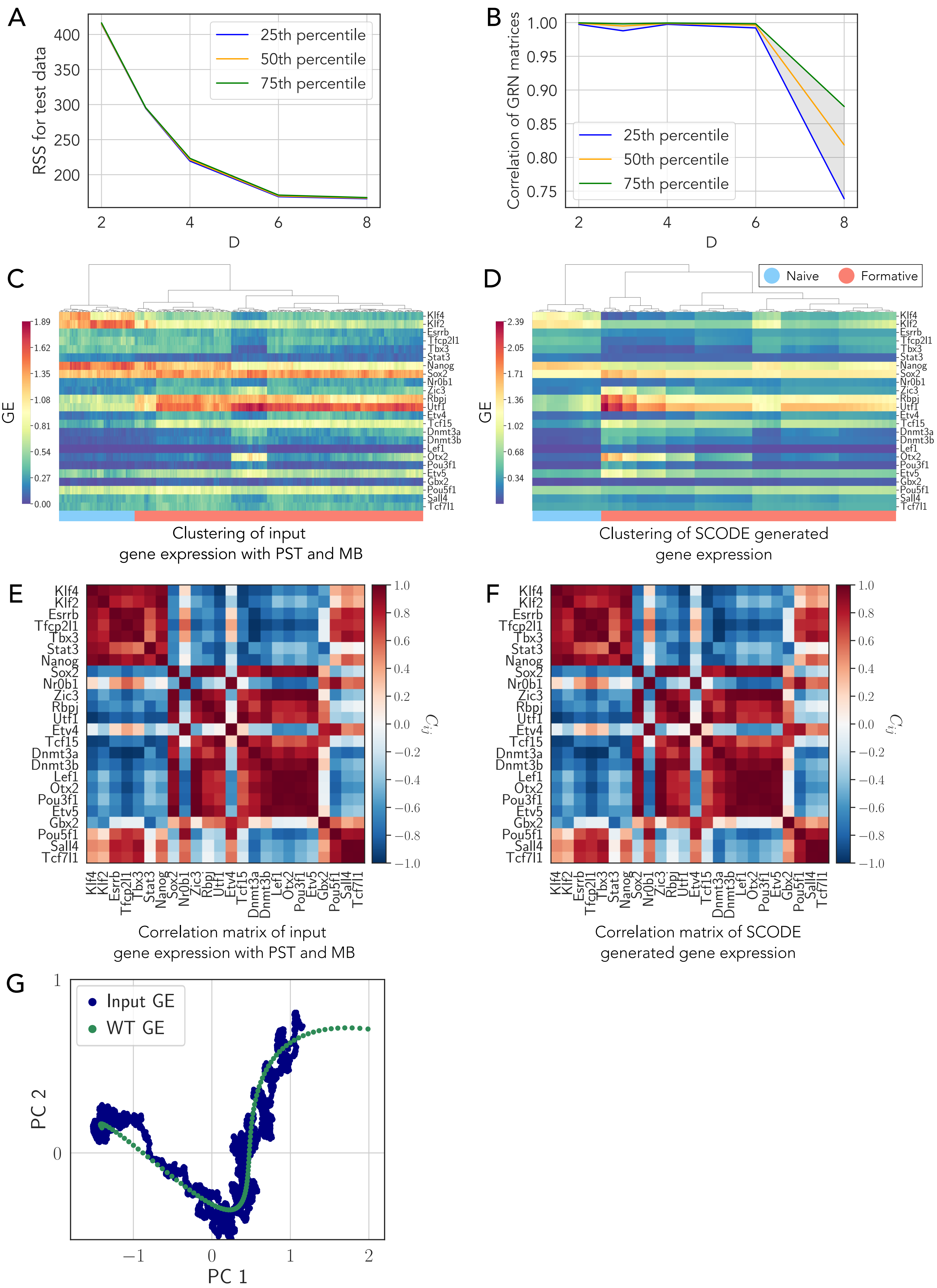

A
RSS for test data
25th percentile
50th percentile
75th percentile
D
B
Correlation of GRN matrices
C
GE
Klf4
Klf2
Esrrb
Tfcp2l1
Tbx3
Stat3
Nanog
Sox2
Nr0b1
Zic3
Rbpj
Utf1
Etv4
Tcf15
Dnmt3a
Dnmt3b
Lef1
Otx2
Pou3f1
Etv5
Gbx2
Pou5f1
Sall4
Tcf7l1
Clustering of input gene expression with PST and MB
D
Naive
Formative
Clustering of SCODE generated gene expression
E
$C_{ij}$
Correlation matrix of input gene expression with PST and MB
F
Correlation matrix of SCODE generated gene expression
G
Input GE
WT GE
PC 1
PC 2

#### Figure S2

A. First, second and third percentiles of the residual sum of squares (RSS) values of the test data for different D (dimensionality of the reduced expression dynamics representation, see Methods for details) values in SCODE analysis.

B. First, second and third percentiles of the correlation values among the optimised GRNs for the top 50 replicates (in ascending order of RSS values) of test data for each D in SCODE analysis.

C. Heatmap with hierarchical clustering of gene expression for the input dataset (scRNA-seq data with LogNorm, pseudotime, PST, and Mini-Bulk, MB). Methodology for visualisation as in Fig. 2D. The dataset has 9547 cells.

D. Heatmap with hierarchical clustering of gene activity of SCODE-generated data. Methodology for visualisation as in Fig. 2D. 100 cells were simulated, as suggested by the authors of SCODE [10].

E. Pearson correlation matrix for the input gene expression dataset.

F. Pearson correlation matrix for SCODE-generated gene expression dataset.

G. Two-dimensional PCA scatter plot representing the gene expression (GE) for the input dataset (scRNA-seq data with LogNorm, PST, and MB) and the SCODE-generated wild-type (WT) data, using the same dimensional reduction approach as in Fig. 2F. Each point corresponds to a single cell. PC1 and PC2 indicate the two dimensions of the PCA space.

### FIGURE S3

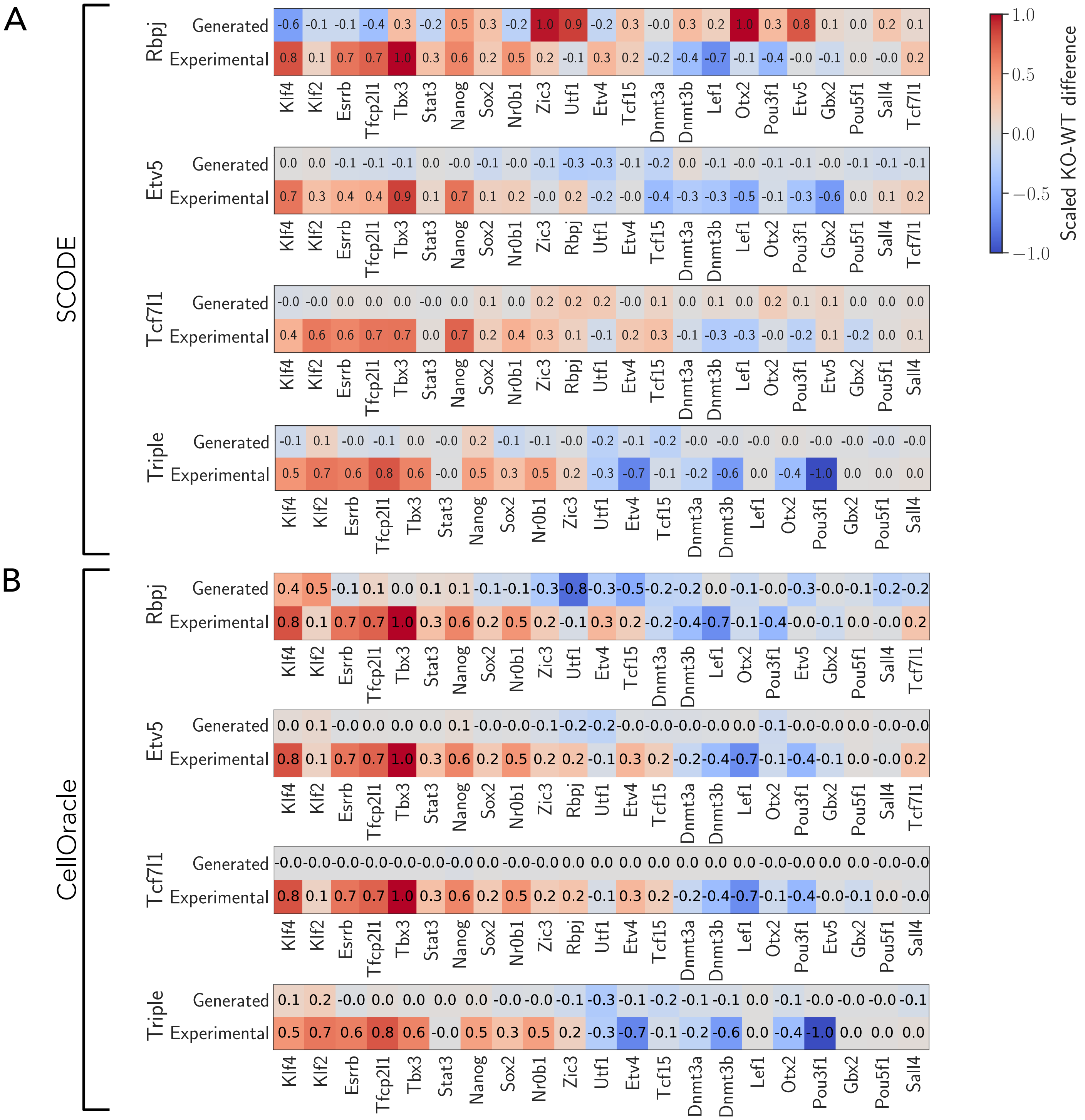
A
SCODE
Rbpj
Etv5
Tcf7l1
Triple
Generated
Experimental
B
CellOracle
Klf4
Klf2
Esrrb
Tfcp2l1
Tbx3
Stat3
Nanog
Sox2
Nr0b1
Zic3
Rbpj
Utf1
Etv4
Tcf15
Dnmt3a
Dnmt3b
Lef1
Otx2
Pou3f1
Etv5
Gbx2
Pou5f1
Sall4
Tcf7l1
Scaled KO-WT difference
1.0
0.5
0.0
−0.5
−1.0

#### Figure S3

A. Scaled KO-WT difference for Rbpj, Etv5, Tcf7l1 and triple KO simulations with SCODE and comparison with experimental scaled KO-WT difference data [30, 29]. For each gene, to compute the simulated scaled KO-WT difference, we measured the scaled difference between the average of the gene expression data in wild-type and knockout conditions. For the experimental data, we scaled the log2FC values measured in [30]. The scaling of all quantities is between $-1$ and $+1$ to facilitate the comparison between different datasets (see Methods for details).

B. Scaled KO-WT difference for Rbpj, Etv5, Tcf7l1 and triple KO simulations with CellOralce and comparison with experimental scaled KO-WT difference data [30, 29]. Methodology for computing the simulated end experimental scaled KO-WT difference as in Fig. S3A (see Methods for details).

### FIGURE S4

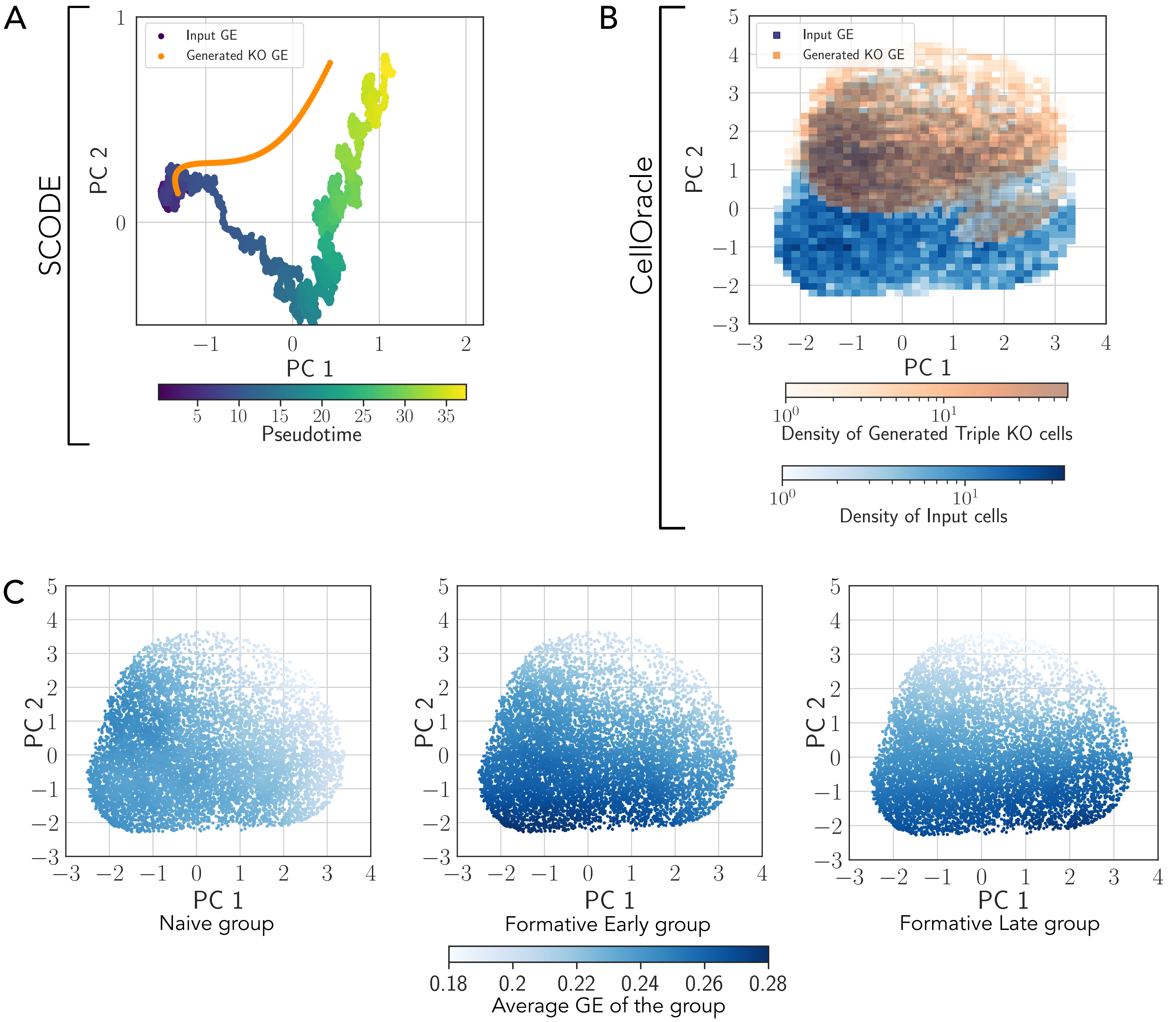

A
SCODE
Input GE
Generated KO GE
PC 2
PC 1
Pseudotime
B
CellOracle
Input GE
Generated KO GE
PC 2
PC 1
Density of Generated Triple KO cells
Density of Input cells
C
PC 2
PC 1
Naive group
Formative Early group
Formative Late group
Average GE of the group

#### Figure S4

A. Two-dimensional PCA scatter plot representing the gene epression, GE, of the input dataset (scRNA-seq data with LogNorm, PST, and MB) and the SCODE-generated triple KO GE. Each point corresponds to a single cell. For the input GE, the colour intensity reflects the pseudotime value of each cell.

B. Two-dimensional PCA scatter plot representing the GE of the input dataset and the CellOracle-generated triple KO GE. Each point corresponds to a single cell. The colour gradient represents the cell density within each square area, with separate scales for the input (blue) and the generated triple KO cells (orange). PC1 and PC2 indicate the two dimensions of the PCA space.

C. Two-dimensional PCA scatter plot representing the GE of the CellOracle input dataset (scRNA-seq data with LogNorm). The colour gradient represents the average gene expression for each group of genes (naive, formative early and formative late, respectively). PC1 and PC2 indicate the two dimensions of the PCA space.

### FIGURE S5

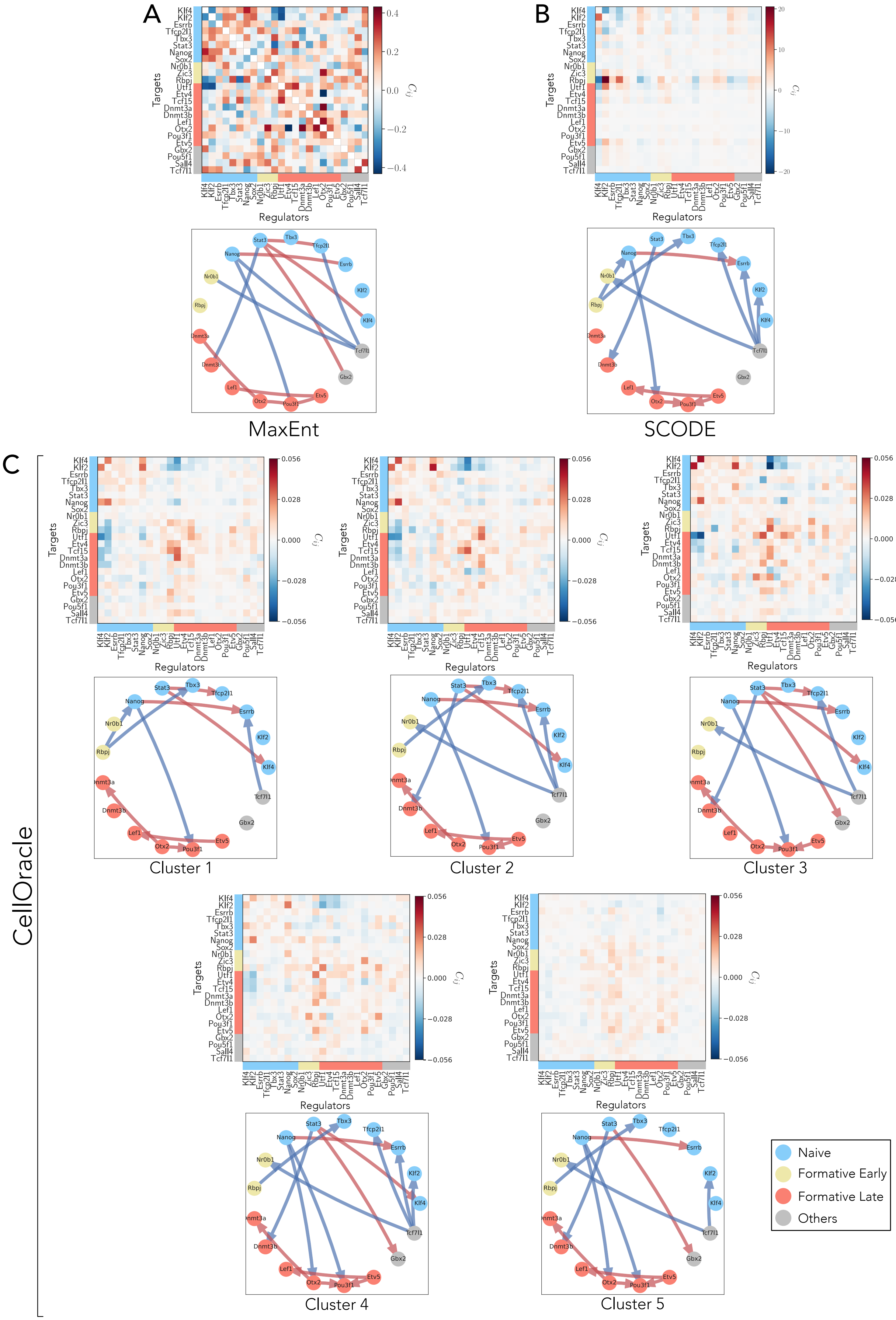

A
B
C
Targets
Regulators
$C_{ij}$
Klf4
Klf2
Esrrb
Tfcp2l1
Tbx3
Stat3
Nanog
Sox2
Nr0b1
Zic3
Rbpj
Utf1
Etv4
Tcf15
Dnmt3a
Dnmt3b
Lef1
Otx2
Pou3f1
Etv5
Gbx2
Pou5f1
Sall4
Tcf7l1
MaxEnt
SCODE
CellOracle
Cluster 1
Cluster 2
Cluster 3
Cluster 4
Cluster 5
Naive
Formative Early
Formative Late
Others

#### Figure S5

A. MaxEnt undirected GRN. Upper panel: interaction matrix computed using the input dataset (scRNA-seq data with LogNorm, PST, and MB), presented in a style analogous to Fig. 4A. Lower panel: GRN of the subset of interactions known from the literature correctly inferred with MaxEnt, presented in a style analogous to Fig. 4C.

B. SCODE GRN. Upper panel: interaction matrix computed using the input dataset (scRNA-seq data with LogNorm, PST, and MB), presented in a style analogous to Fig. 4A. Lower panel: GRN of the subset of interactions known from the literature correctly inferred with SCODE, presented in a style analogous to Fig. 4C.

C. CellOracle GRNs computed using the input dataset (scRNA-seq data with LogNorm) for each cluster within the dataset. Upper panels: CellOracle interaction matrices for each cluster, presented in a style analogous to Fig. 4A. Lower panels: GRN of the subset of interactions known from the literature correctly inferred with CellOracle, presented in a style analogous to Fig. 4C.

### FIGURE S6

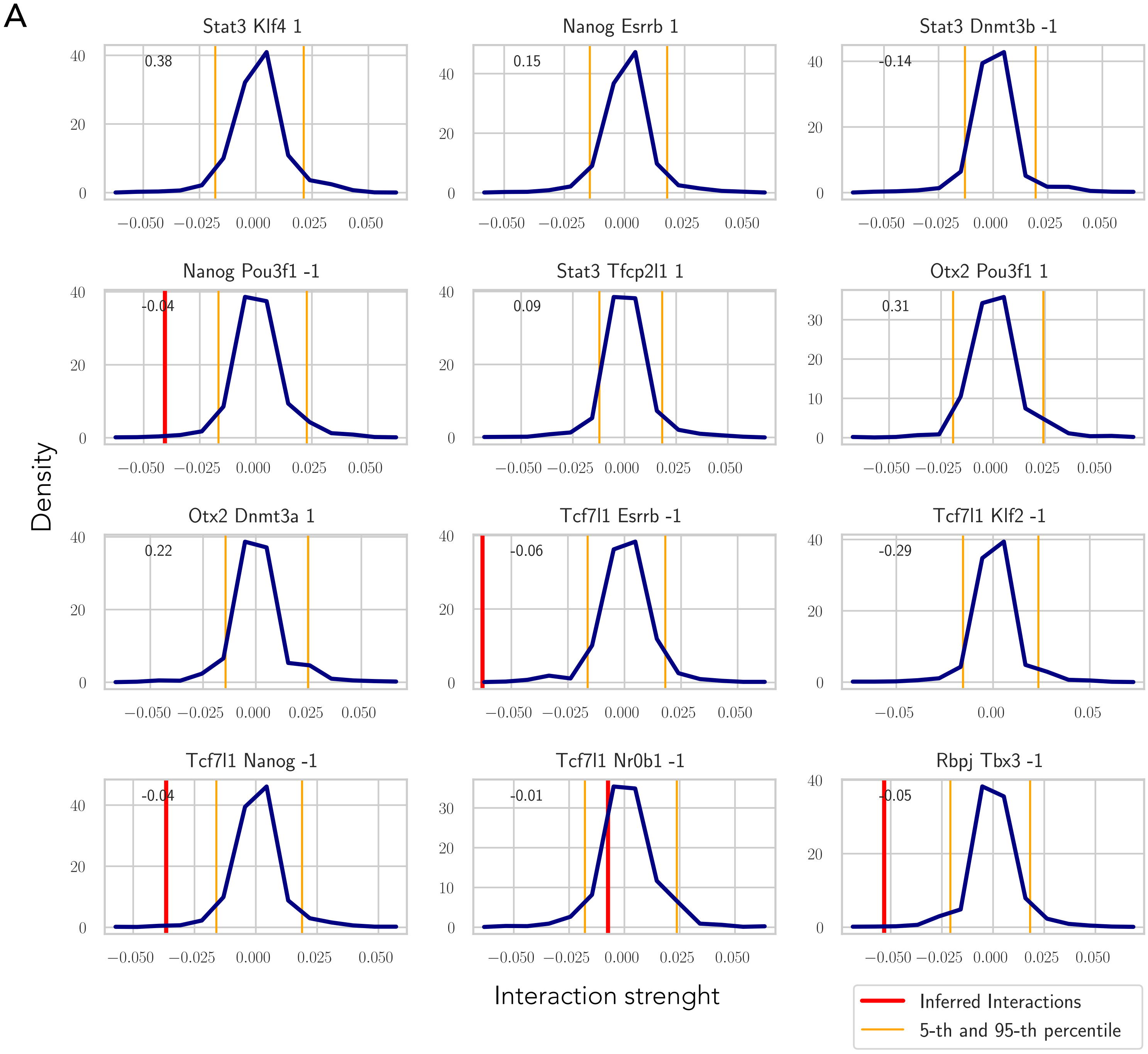
A
Stat3 Klf4 1
0.38
Nanog Esrrb 1
0.15
Stat3 Dnmt3b -1
-0.14
Nanog Pou3f1 -1
-0.04
Stat3 Tfcp2l1 1
0.09
Otx2 Pou3f1 1
0.31
Otx2 Dnmt3a 1
0.22
Tcf7l1 Esrrb -1
-0.06
Tcf7l1 Klf2 -1
-0.29
Tcf7l1 Nanog -1
-0.04
Tcf7l1 Nr0b1 -1
-0.01
Rbpj Tbx3 -1
-0.05
Density
Interaction strenght
Inferred Interactions
5-th and 95-th percentile

#### Figure S6

A. Distribution of interaction values for selected pairs of genes from 2500 GRNs. These networks are inferred with IGNITE using 50 shuffled datasets as input. These datasets are randomly shuffled versions of the input dataset (scRNA-seq data with LogNorm, PST, and MB). Then for each of these datasets 50 sets of hyperparameters were used (see Methods for details). Each panel illustrates the distribution of specific gene pair interaction values across the IGNITE-derived GRNs (with $N_{bins} = 15$). The red lines indicate the interaction values from the IGNITE GRN with the lowest CMD, while the yellow lines delineate the fifth and 95th percentile confidence intervals, providing a basis for evaluating the significance of each interaction.
